# Galectin-1 identifies a unique subpopulation of highly invasive glioblastoma cells and enables their migration

**DOI:** 10.64898/2026.06.15.732361

**Authors:** Talia Sanazzaro, Michael Cotner, Neha Arvinth, Amy Brock, Stephanie K Seidlits

**Affiliations:** Department of Biomedical Engineering, The University of Texas, Austin, Texas, USA

**Author notes:** **Corresponding Author:** Stephaine K Seidlits.

## Abstract

Glioblastoma (GBM), the most common primary brain tumor, is characterized by extensive infiltration into surrounding brain tissue. GBM tumors exhibit substantial intratumoral heterogeneity making it difficult to identify and target invasive cell subpopulations. Here, we use an *in vitro* model of the mechanical transitions at the tumor-brain interface to isolate highly invasive GBM cells from populations derived from unique patient tumors. Using single-cell RNA sequencing combined with cell barcodes we were able to trace distinct cell lineages during migration and identify an invasive subpopulation which preferentially migrates across stiffness interfaces. This mechanically sensitive invasive subpopulation exhibits a transcriptomic profile characterized by overexpression of galectin-1, a β-galactoside binding protein. Our findings reveal that galectin-1 expression level positively correlates with invasion rate and inhibition of galectin-1 binding to cell surface glycoproteins prevents migration. This study identifies galectin-1 expression is a characteristic of invading GBM cells in mechanically defined interface model where galectin-1 may serve as an important maker to identify and target invasive GBM cells.

## INTRODUCTION

Glioblastoma (GBM), arise *de novo* as a grade IV tumor and is the most common primary brain tumor.^2^ GBM prognosis is poor, with a median survival time of 15 months and a 5-year survival of less than 7%.^3^ Despite aggressive treatments, nearly all patients experience tumor recurrence.^4^ GBM tumors are highly invasive, where cancer cells invade from the tumor periphery to adjacent brain tissue. These invading cells are virtually impossible to resect surgically and likely seed recurrent tumors.^5^ Thus, therapies that specifically target invasive GBM cells are of great interest. In this study, we set out to characterize the transcriptomes of invasive GBM populations and identify unique, targetable features.

GBM tumors are highly heterogeneous, with significant transcriptomic differences observed between cells at the invasive edge and the tumor bulk.^6, 7^ For IDH-WT tumors, malignant GBM cells have been characterized by 4 transcriptomic states; astrocyte-like (AC), oligodendrocyte-progenitor-like (OPC), neural progenitor-like (NPC), and mesenchymal-like (MES).^8^ All 4 cellular states are present throughout primary patient tumors but at varying proportions and it has been demonstrated that cells from any of these 4 subtypes have the capacity to invade from the tumor edge.^9, 10^ Conventional scRNAseq is widely leveraged to investigate the transcriptomes of cells and identifies distinct subpopulations in GBM and other contexts.^11^ However, these snapshot measurements cannot distinguish whether cell state alterations result from individual cells undergoing a change in transcriptomic state or from a shift in the abundance of different subsets of cells. To distinguish this, researchers have developed technologies that enable tracking of individual clonal subpopulations following experimental perturbation, such as the ClonMapper technology used in this study.^12, 13^ By combining cell barcodes and scRNAseq, it becomes possible to analyze the transcriptomic expression profiles of specific subpopulations, identify subpopulation-specific responses, and begin to understand how intratumoral heterogeneity contributes to disease progression.

Invasive GBM cells must migrate through the brain extracellular matrix (ECM), which is enriched in large glycosylated biomolecules with the glycosaminoglycan hyaluronic acid (HA) in abundance.^14, 15^ GBM cells produce various ECM components at abnormal rates, with increased deposition of HA, tenascin-C, fibronectin, and fibrillar collagens.^16–19^ Increased ECM deposition in tumors provides more sites for cell-matrix adhesion and may facilitate GBM cell migration.^15, 20^ Furthermore, increased ECM deposition results in regional alterations in tissue mechanics where the tumor core is consistently 10-fold stiffer (compressive Young’s modulus) than the adjacent peri-tumoral tissue—at least in orthotopically xenografted patient-derived GBM tumors in mice.^21^ Mechanotransduction has been studied in a variety of biological processes including the initiation and progression of GBM.^22, 23^ Previous studies have suggested that the change in mechanical modulus from stiffer tumor tissue to softer peritumoral brain tissue can drive migration and oncogenic gene expression.^24, 25^ For example, previous reports have found that GBM cells cultured in soft matrices migrate more readily than those in stiffer matrices.^21, 26, 27^ Here, we take this concept further, investigating how mechanical interfaces affect heterogeneous GBM cell populations in ways that promote diffuse invasion.

In this study we employed a 3D culture model of the mechanical interface encountered by cells at the tumor edge. Cells were cultured in ECM-derived hydrogels, polymerized to mimic the stiffnesses of the peritumoral tissue and tumor edge, to model the stiffness interface that occurs naturally in GBM tumors. We then investigated the transcriptomic changes in patient-derived GBM cells that migrate across this mechanical interface. Using scRNAseq combined with clonal lineage tracing of barcoded GBM cell populations, we identified an invasive subpopulation which preferentially migrated across mechanical interfaces and the unique transcriptomic identity of this invasive subpopulation. Galectin-1, a β-galactoside binding protein, was found to be a signature marker of a unique mechano-responsive clonal subpopulation. We confirm that galectin-1 protein expression is a feature of xenografted GBM cells invading along blood vessels in a mouse model. Moreover, galectin-1 was shown to be required for GBM cell migration *in vitro* in patient-derived GBM models.

## RESULTS

### GBM cells primarily migrate across stiffness interface into soft matrices

Two hydrogels—one approximating the stiffness of GBM tumor tissue and the other that of peritumoral brain tissue—were positioned in direct contact with each other to model the mechanical transition at the invasive edge of GBM tumors. Shear storage moduli (G’) were confirmed to be 1000 Pa +/− 100 for the stiff hydrogel matrix and 100 Pa +/− 50 for the soft hydrogel matrix, mimicking the stiffnesses of GBM tumor tissue and peritumoral brain tissue, respectively (**Figure 1A**).^21^ Three interface conditions were evaluated as follows: soft → soft, stiff → soft, soft → stiff, with cells from the patient-derived GBM cell line, HK177, 3D cultured **only** in the first hydrogel. The rate of invasion across the mechanical interface was calculated based on area coverage (i.e., confluence) in the adjacent, cell-free gel over 8 days. While GBM cells initially seeded in the stiffer matrix rapidly invaded into the softer matrix (stiff → soft condition), minimal invasion was observed in the soft → stiff condition (**Figure 1B/C**). Invasion was also observed in the soft → soft condition. No migration is observed without the presence of a soft matrix, as we have previously reported.^21^

**Figure 1:**
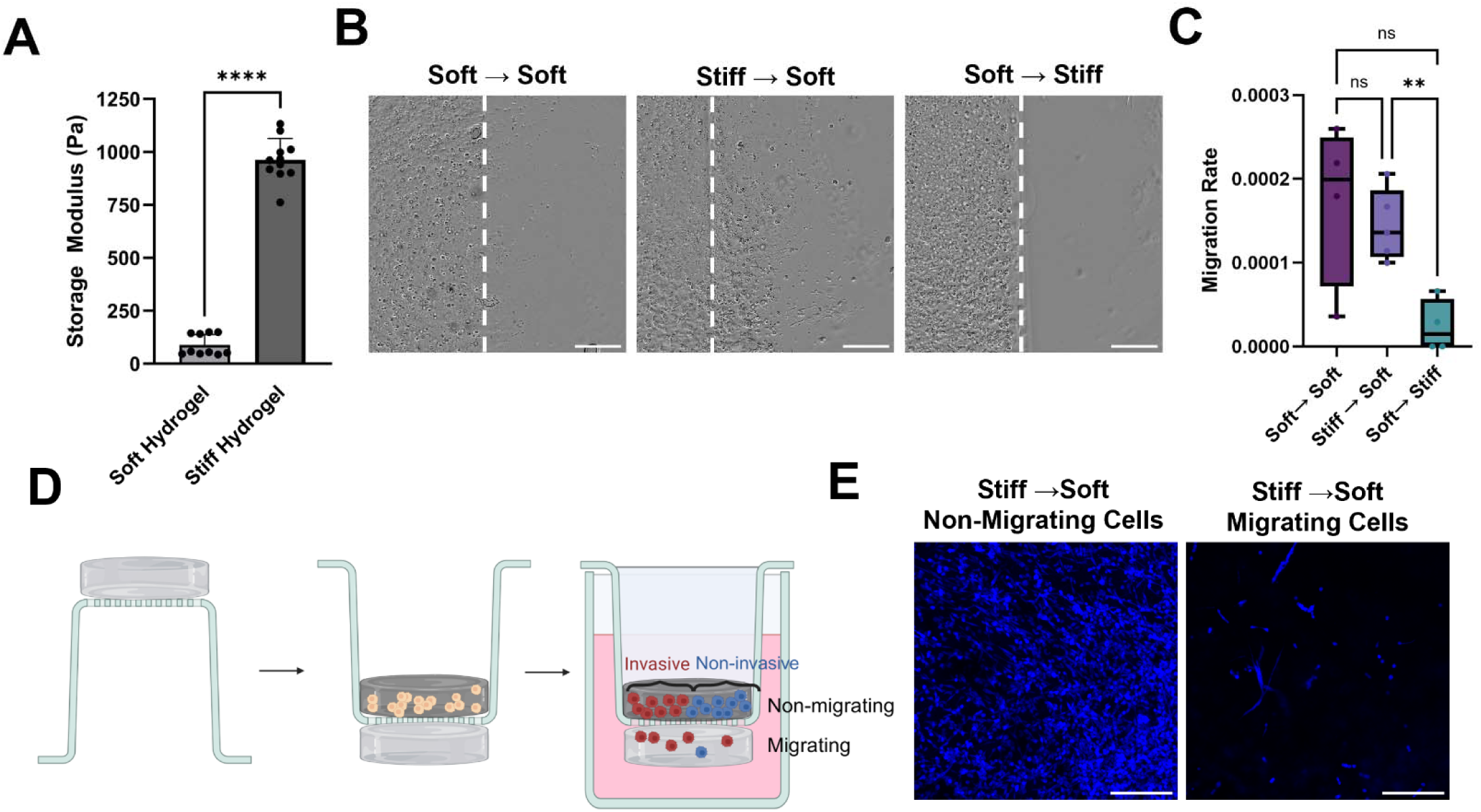
Interface hydrogel model to mimic mechanical transition seen at tumor-brain interface. **A.** 10-fold difference in storage modulus (G’) between soft and stiff hydrogels. Unpaired two-sided t-test, n=10, *\*\*\*\* p =* 1.13*10^−13^. Error bars represent mean with standard deviation. **B.** Representative images of cell line HK177 migration in the interface hydrogel conditions after 8 days. Scale bar = 500 μm **C**. Cells have a higher migration rate across interfaces into soft matrices over 8 days. Migration rate is defined as %confluence to the right of the interface due to migrating cells per hour. One-way ANOVA with Dunnett T3 post hoc test, n=5 technical replicates, ** *p =* 0.005, ns = no significance. Error bars represent min to max values. **D.** Schematic of interface hydrogel model for separating cells for scRNAseq and a visual for the naming conventions used. Figure created in BioRender. Sanazzaro, T. (2026) https://BioRender.com/jlzgn51. **E.** Maximum z-projection confocal images of cells in stiff → soft interface hydrogel model at day 14. Scale bar = 200 μm.

ClonMapper cell barcoding was used to identify whether mechanically sensitive invasive subpopulations exist that preferentially migrate across the stiff → soft mechanical interface.^12^ A patient-derived GBM cell line HK177^28^ was transduced with genetically integrated cellular barcodes.^12^ Barcoded HK177 cells had high clonal diversity with roughly 300 unique barcode sequences identified in scRNAseq with the most abundant barcode sequence comprising roughly 10% of the population (**Figure S1**).

ScRNAseq was used to investigate the effects of stiffness on the transcriptome of migrating and non-migrating cells in the mechanical interface hydrogel cultures. Two GBM patient-derived lines, barcoded HK177 cells and GS54, a non-barcoded line, were used in these studies.^28, 29^ For scRNAseq experiments, Corning HTS Transwell plates were used to separate cells that had actively migrated from the original seeding position from those cells that remained. Cells were encapsulated into the stiff, tumor-like hydrogel at the top of the Transwell and interfaced with the soft, brain-like hydrogel on the bottom of the Transwell (**Figure 1D**). A soft → soft interface hydrogel was used as a control, as there was no change in mechanical modulus. Although our previous experiment revealed minimal invasion in the soft → soft condition, we sought to investigate both the features of mechanically driven invasive cells preconditioned in a stiff microenvironment (stiff → soft), and the transcriptomic features of cells which migrate and do not migrate, regardless of matrix stiffness, to better elucidate the how matrix mechanics affect the transcriptome of GBM cells in the context of tumor cell invasion.

Cells were cultured in the interface hydrogels for 2 weeks. **Figure 1E** provides an example (stiff → soft condition) of the relative densities of the HK177 cells on the original (stiff) side of the interface, which we will refer to as the “non-migrating cells”, and the opposite (soft) side, which we will refer to as the “migrating cells” after 14 days in culture. At this point, the hydrogels were separated and cells were extracted from each side. Four experimental groups were collected for scRNAseq: soft → soft migrating, soft → soft non-migrating, stiff → soft migrating, stiff → soft non-migrating. Following alignment, initial quality control and filtering, 4943 cells and 9619 genes from GS54 and 3850 cells and 8873 genes from HK177 were analyzed.

### GBM samples are highly heterogeneous, and transcriptomic profile is correlated with migration status

To initially visualize the transcriptomic data, cells were projected onto a two-dimensional space through uniform manifold approximation and projection (UMAP) and labeled by experimental group. Interestingly, HK177 clustered primarily by migration status, rather than by stiffness condition, while GS54 clustered by migration status and stiffness condition. (**Figure 2A/E**). To evaluate the heterogeneity within each GBM model, analysis was conducted independently for HK177 and GS54.

**Figure 2:**
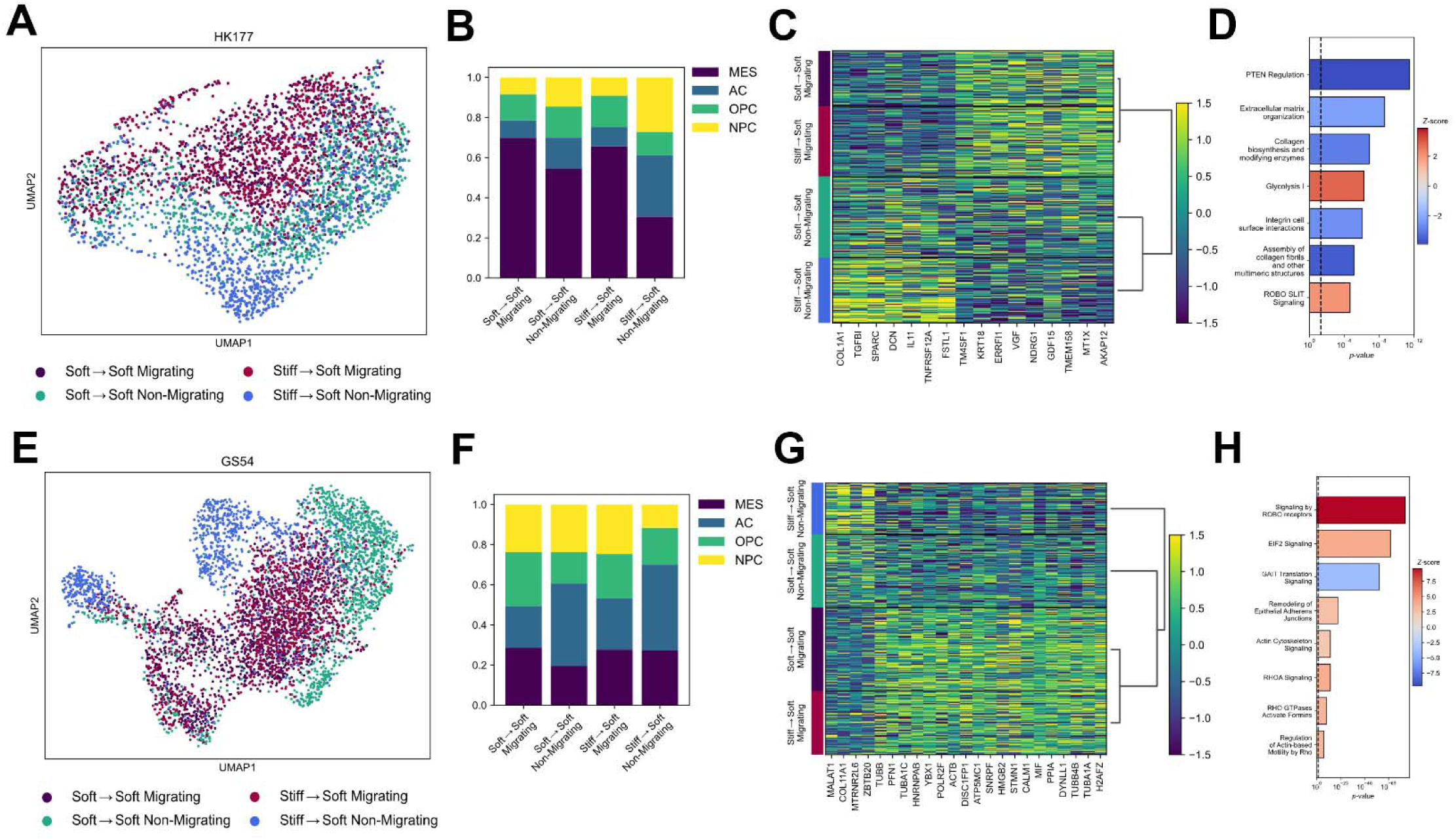
GBM samples display heterogeneity in tumor-interface model. **A.** UMAP of HK177 **B**. Malig ant cell type characterization separated by sample condition of HK177. A two-sided unpaired t-test was used to calculate differences in MES, AC, OPC, and NPC score per cell between conditions (n = 786 soft → soft migrating, n = 1143 soft → soft non-migrating, n = 998 stiff → soft migrating, n = 923 stiff → soft non-migrating). False discovery rate control was performed using Benjamini-Hochberg-adjusted p-values. Statistical data can be accessed in the supplemental data. **C.** Heatmap of most differentially expressed genes identified from MAST between migrating (n = 1784) and non-migrating (n = 2066) cells in HK177 (*p* < 5*10^−5^, log_2_FC > 0.75). False discovery rate control was performed using Benjamini-Hochberg-adjusted p-values. **D.** IPA identifies the differentially regulated pathways between non-migrating and migrating cells in HK177. Positive z-score (red) corresponds to an upregulation of a pathway in the migrating cells, while negative z-score (blue) corresponds to pathways upregulated in the non-migrating cells. **E.** UMAP of GS54. **F**. Malignant cell type characterization separated by sample condition of GS54. A two-sided unpaired t-test was used to calculate differences in MES, AC, OPC, and NPC score per cell between conditions (n = 1519 soft → soft migrating, n =1331 soft → soft non-migrating, n = 1157 stiff → soft migrating, n = 936 stiff → soft non-migrating). False discovery rate control was performed using Benjamini-Hochberg-adjusted p-values. **G.** Heatmap of most differentially expressed genes identified from MAST between migrating (n = 2676) and non-migrating (n = 2267) cells in GS54 (*p* < 5*10^−5^, log_2_FC > 0.6). False discovery rate control was performed using Benjamini-Hochberg-adjusted p-values. **H.** IPA identifies the differentially regulated pathways between non-migrating and migrating cells in HK177. Positive z-score (red) corresponds to an upregulation of a pathway in the migrating cells, while negative z-score (blue) corresponds to pathways upregulated in the non-migrating cells.

The abundance of the four GBM malignant cell types (MES-like, NPC-like, OPC-like and AC-like) of each experimental group was analyzed using datasets identified by Neftel et al.^8^ The gene used to determine the cell type analysis are available in the supplemental data. Both patient lines display heterogeneity with expression of all malignant cell states. Both HK177 and GS54 cell populations had an initial TCGA (The Cancer Genome Atlas) classification of mesenchymal, based on HK177 containing an *EGFRvIII* mutation,^28^ and GS54 containing mutations in *PTEN*, *NF1* and a deletion of *CDKN2A*.^29^ For HK177 which was comprised of a large fraction of mesenchymal-like (MES) cells, stiffness affected the cell state (**Figure 2B**). There was a 2-fold increase in MES-like cells collected from soft matrices (soft → soft migrating: *p* = 7.7*10^−61^, soft → soft non-migrating: *p* = 3.3*10^−39^, stiff → soft migrating: *p* = 1.5*10^−58^) compared to the cells collected from the stiff matrix (stiff → soft non-migrating). While the fraction of AC and NPC-like cells was increased in cells collected from the stiff matrix, with a 2-fold increase in cells from stiff → soft non-migrating, compared to soft → soft migrating (AC: *p* = 2*10^−21^, NPC: *p* = 3.3*10^−28^), soft → soft non-migrating (AC: *p* = 3.3*10^−8^, NPC: *p* = 4.4*10^−11^), stiff → soft migrating (AC: *p* = 1.8*10^−18^, NPC: *p* = 3.5*10^−25^). The fraction of OPC-like cells was similar across all conditions for HK177 cells.

To understand the differences between migrating and non-migrating cells, differential gene expression analysis (DGEA) was conducted to compare non-migrating (cells that remained in the top hydrogel: soft → soft non-migrating, stiff → soft non-migrating) and migrating (cells that migrated into the bottom hydrogel: soft → soft migrating, stiff → soft migrating) cells. HK177 cells that did not migrate upregulated (*p* < 5*10^−5^, log_2_FC > 0.75) the ECM-related genes *COL1A1, TGF*β*1, SPARC,* and *DCN* (**Figure 2C**). Ingenuity Pathway Analysis (IPA) was used to determine differentially regulated pathways. Cells which did not migrate upregulated pathways involved with the ECM including “*Extracellular matrix organization*”, “*Collagen biosynthesis and modifying enzymes”,* “*Assembly of collagen fibrils and other multimeric structures*”, and “*Integrin cell surface interactions*” (**Figure 2D**). Migrating cells highly upregulated (*p* < 5*10^6^, log_2_FC > 0.75) several migration-associated genes, including *AKAP12, TMEM158, TM4SF1, KRT18, STC1,* and *HMGA1* (**Figure 2C**). Corresponding to an upregulation of “ROBO SLIT Signaling” and “*Glycolysis I*” pathway which relates to metabolic functions of the migrating cells (**Figure 2D**).

For GS54 cells in both stiffness interface conditions, cells that did not migrate (stiff → soft non-migrating and soft → soft non-migrating) had an increase in AC-like cells compared to the cells that did migrate (soft → soft migrating: *p* = 5.2*10^−64^ and stiff → soft migrating: *p* = 3.3*10^−48^) (**Figure 2F**). The statistical analysis for all comparisons of both cell lines can be found in the supplemental data. In the GS54 cell line the non-migrating GS54 cells upregulated (*p* < 5*10^−5^, log_2_FC > 0.6) the ECM-related gene, *COL11A1* while the migrating GS54 cells upregulated cytoskeleton-related genes, specifically *TUBA1A, TUBB4B, TUBB, TUBA1C, STMN1, DISC1, HNRNPAB, ACTB,* and *PFN1* (**Figure 2G**). Together, these changes correspond to an upregulation of “*Signaling by ROBO receptors*”, “*EIF2 signaling*” and “*Actin cytoskeleton signaling”* pathways as well as an upregulation in various Rho signaling pathways. The non-migrating cells had enriched expression of transcripts related to “*GAIT Translation Signaling Pathway*” (**Figure 2H**).

### Invasive subpopulations exist within HK177 which preferentially migrate across interface

If GBM cell populations display heterogeneous levels of invasiveness, highly invasive cell subpopulations would be enriched in the migrating cell groups. To this end, we leveraged the ClonMapper cell barcoding technology to track the invasiveness of mechanically driven clonal cell subpopulations of HK177 during migration across the stiffness interface (stiff→ soft) with integrated DNA barcodes. For each barcode, a binomial test was used to determine if the number of cells with that barcode was significantly higher than random chance in the migrating or non-migrating groups of the stiff → soft condition. There were 10 barcodes which were significantly overrepresented in the non-migrating group and 5 barcodes which were significantly overrepresented in the migrating group (adjusted *p* < 0.05), assigned as non-invasive and invasive subpopulations, respectively (**Figure 3A**).

**Figure 3:**
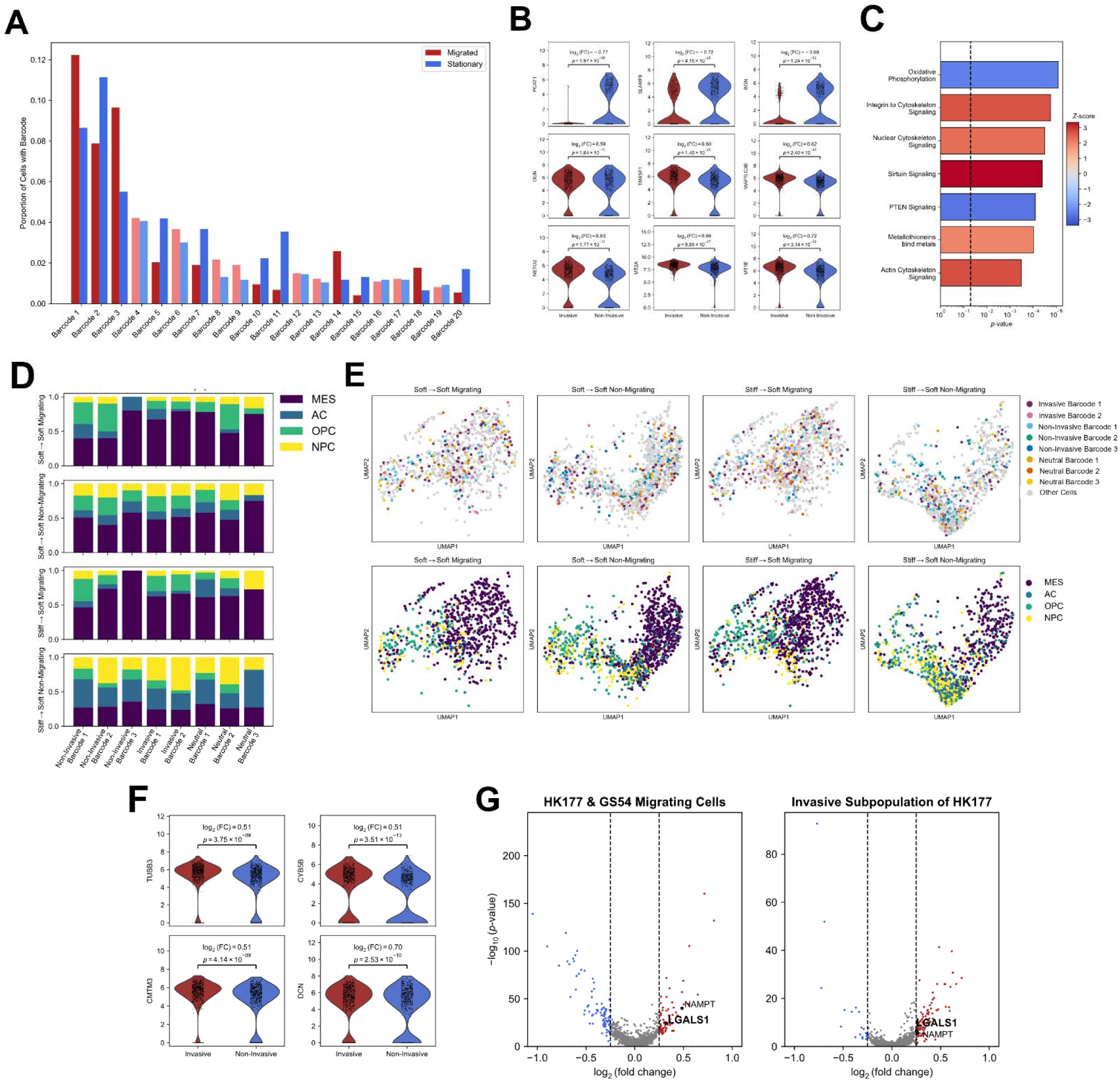
Invasive subpopulations exist which predominantly migrate at the interface. **A**. A binomial test determined that significantly significant invasive and non-invasive lineages exist (*p* < 0.05). Light colored bars indicate barcode abundance is not significantly different in either gel. **B**. Differentially expressed genes identified from MAST between the invasive (n = 641) and non-invasive (n = 655) subpopulations (*p* < 5*10^−5^, log_2_FC > 0.6). False discovery rate control was performed using Benjamini-Hochberg-adjusted p-values. Adjusted p-values for each gene are displayed on the graph. **C.** Differentially expressed pathways identified by IPA between the invasive and non-invasive subpopulations. Positive z-score (red) corresponds to an upregulation of a pathway in the migrating cells, while negative z-score (blue) corresponds to pathways upregulated in the non-migrating cells. **D.** The composition of malignant GBM cell states within individual clonal subpopulations. **E**. UMAP demonstrates that cells are more closely colocalize by malignant cell state (bottom) rather than clonal subpopulation (top). **F.** Differentially expressed genes identified from MAST between the invasive (n = 641) and non-invasive (n = 655) subpopulations that can be attributed to stiffness (*p* < 5*10^−5^, log_2_FC > 0.5). False discovery rate control was performed using Benjamini-Hochberg-adjusted p-values. Adjusted p-values for each gene are displayed on the graph. **G.** Expression of LGALS1 identifies migrating cells in HK177 and GS54 (left) and the invasive subpopulation of HK177 (right).

To determine genes that are correlated with an enhanced invasive phenotype, the invasive subpopulation (all cells with an invasive barcode sequence) and the non-invasive subpopulation (all cells with an invasive barcode sequence) from HK177 were compared. Invasive and non-invasive cells were combined from all conditions, and DGEA was conducted with matrix stiffness and migration modeled as a random effect. These differentially expressed genes can be used to mark a “pre-invasive” phenotype in the HK177 line. Notably, overexpression (*p* < 5*10^−5^, log_2_FC > 0.6) of several genes from the metallothionein family, *MT1E* and *MT2A* and the metastasis-related genes *NETO2* and *TM4SF1* were correlated with an invasive phenotype, while the ECM proteoglycan *BGN* and cancer-progression associated genes *PCAT1, SLAMF9* and *PVT1* were upregulated in the non-invasive subpopulation (**Figure 3B**). Pathways associated with the pre-invasive phenotype (*p* < 5*10^−3^, z-score > 2) included “*Metallothionein bind metals*” and cell migration-associated pathways including “*Integrin to cytoskeleton signaling,*” “*Actin cytoskeleton signaling,*” and “*Nuclear cytoskeleton signaling*” while upregulation of “*PTEN signaling*” and “*Oxidative Phosphorylation*” correlated to the non-invasive subpopulations (**Figure 3C**). There were no observed significant differences in the cell cycle phase between the subpopulations further suggesting that the identified genes are not driven by cell-cycle-dependent differences in gene expression (**Figure S2A**).

The cell states of the invasive and non-invasive subpopulation were similarly comprised of the 4 cell types (**Figure S2B**), so we sought to investigate how cell state shifts within clonal derived cell populations. The statistical analysis can be accessed in the supplemental data. For the most abundant barcodes of HK177 (barcodes with a minimum of 10 cells in each group), the composition of the 4 GBM cell states was calculated (**Figure 3D**). All four cell types were represented across all these most abundant clonal subpopulations, rather than subpopulations being mostly comprised of single cell types. This result indicates that between the initial barcoding of single cells and experimental endpoint, cells within a single clonal subpopulation quickly diversified to represent all four GBM subtypes, despite being relatively recent descendants of the same cell. On a UMAP scatter plot, cells colocalize by cell type rather than clonal identity, directly contrasting what is typically observed in similar cell barcoding experiments (**Figure 3E**).^12, 30^ Across all clonal subpopulations, matrix stiffness appeared to have the greatest effect on the cell state where a reduction in MES-like cells was observed coinciding with an increase in AC and NPC-like cells. The cutoff number of cells for each barcode in each condition was 10, so these observed trends are limited by the low number of cells.

We hypothesized that stiffness has different effects on the transcriptome of the invasive and non-invasive subpopulations. In this analysis, invasive and non-invasive subpopulations were compared by DGEA, conducted separately for stiff and soft matrices to analyze how the differences between the two subpopulations change in response to matrix stiffness. 4 genes were identified as differentially expressed (*p* < 5*10^−5^, log_2_FC > 0.5) in the stiff matrix that were not in the soft matrix, including *TUBB3*, *CYB5B*, C*MTM3* and *DCN*, representing markers of an invasive subpopulation induced by a stiff matrix (**Figure 3F**). These results indicate that microenvironmental stiffness can play a role in functional intratumoral heterogeneity.

To identify transcriptomic markers of an invasive cell subpopulation that were also enriched in migrating cells, the upregulated genes from the migrating cells (stiff → soft and soft → soft conditions the set of genes that distinguished the invasive subpopulation from the non-invasive subpopulation in HK177 and which distinguished the migrating and non-migrating cells of both HK177 and GS54 were compared. Here, the two cell lines were combined, and the cell line was modeled as a random effect. Two genes, *LGALS1* and *NAMPT*, were identified as upregulated by the migrating cells of HK177 and GS54 and upregulated by the invasive subpopulation of HK177 (**Figure 3G**). To confirm that these genes represent a unique subpopulation of HK177, MAST differential gene expression analysis was re-evaluated controlling for cell phase and MES cell score, and both genes still distinguished the two subpopulations (**Figure S2C**). There was no observed significant difference in the expression of the hypoxia marker *HIF1A* between the subpopulations, nor HIF1a protein expression in the stiff or soft hydrogels as previously reported at this timescale.^21^

### Substrate stiffness affects the transcriptome of migrating GBM cells

To evaluate the shared effects of matrix stiffness, the two cell lines were combined, and gene expression was compared between the soft→ soft non-migrating cells and the stiff → soft non-migrating cells using MAST DGEA. In these groups, cells did not migrate across the interface at the time of collection and instead remained in a single matrix stiffness. Cells collected from the stiff hydrogel *p* < 5*10^−5^, log_2_FC > 0.7) more highly expressed ECM-related gene *SPARC, VIM,* and *ATF3* and the proto-oncogenes *JUN* and *FOS* compared to the soft → soft non-migrating cells which more highly expressed *DDIT4* and *VEGFA* (**Figure 4A**). Pathway analysis revealed that the ECM related pathways “Actin Cytoskeleton Signaling”, “Regulation of Actin-based Motility by Rho”, “Asparagine N-linked glycosylation” were upregulated by the non-migrating cells in the stiff matrix (**Figure 4B**).

**Figure 4:**
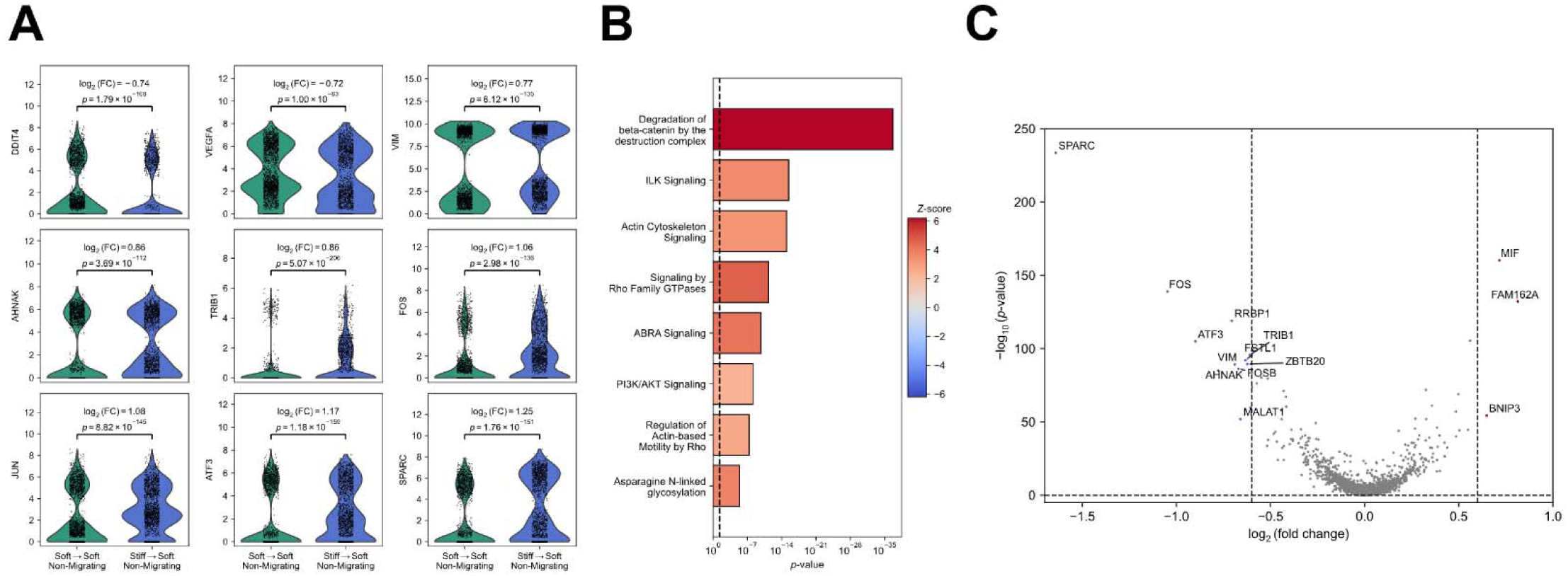
Matrix stiffness affects the transcriptomic expression of GBM cells. **A.** Shared differentially expressed genes identified from MAST between stiff → soft non-migrating (n = 1859) and stiff → soft non-migrating (n = 2474) cells (*p* < 5*10^−5^, log_2_FC > 0.7). False discovery rate control was performed using Benjamini-Hochberg-adjusted p-values. Adjusted p-values for each gene are displayed on the graph **B.** Pathway analysis identified by IPA between stiff → soft non-migratory and stiff → soft non-migratory cells. Positive z-score (red) corresponds to pathway upregulation in cells from the stiff matrix. **C**. Volcano plot of differentially expressed genes between stiff → soft migrating (n = 2155) and stiff → soft non-migrating (n = 1859) cells (*p* < 5*10^−5^, log_2_FC > 0.6). False discovery rate control was performed using Benjamini-Hochberg-adjusted p-values Labeled genes correspond to stiffness-dependent transcriptomic differences migratory cells. on the migratory cells.

To evaluate the effects of matrix stiffness on migrating cells in both cell lines, the gene expression profile between the stiff→ soft migrating cells and the stiff → soft non-migrating cells was compared. To isolate these stiffness dependent effects on migration genes were removed which were differentially expressed (*p* < 5*10^−5^, log_2_FC > 0.5) between the soft → soft migrating and soft → soft non-migrating cells, based on the assumption that these genes correspond to stiffness-independent effects on migration. Similar genes to the previous analysis arose as the non-migrating cells more highly expressed (*p* < 5*10^−5^, log_2_FC > 0.6) the ECM related genes *SPARC, VIM,* and *ATF3* and the proto-oncogenes *JUN, FOS,* and *FOSB* compared to the migrating cells. The migrating cells expressed *MIF* and *FAM162A* when migrating from a stiff to soft matrix (**Figure 4C**).

Persistent stiffness-induced changes on cells that migrated were evaluated by comparing cells that had migrated from the stiff → soft gel and those that had migrated from the soft → soft gel (*p* < 5*10^−5^, log_2_FC > 0.2). Interestingly, there were no differentially expressed genes suggesting that while stiffness may drive the differences in transcriptomic expression between migrating and non-migrating cells, once cells migrate these changes are not permanent since migrating cells are transcriptomically distinct from non-migrating cells.

### Galectin-1 protein is expressed in stiffness interface model and in overexpressed in *in vivo* tumors

Recent literature has shown expression of galectin-1 (*LGALS1*) is enhanced as tumors progress and that overexpression is correlated to worse patient outcomes in various cancers such as breast,^31^ ovarian,^32^ lung,^33^ and colorectal cancer.^34^ Clinical data from the TCGA demonstrated a significantly higher expression of *LGALS1* in GBM compared to non-tumor brain tissue (*p <* 0.0001) (**Figure 5A**).^35, 36^ Additionally, increasing *LGALS1* expression corresponded to a lower survival in GBM patients (**Figure 5B**). Furthermore, spatial data from the IVY GAP dataset demonstrates enhanced expression of LGALS1 is highest along microvascular in GBM tumors (**Figure 5C**). Microvasculature provides a path for invasive GBM cells to exit the primary tumor, allowing GBM cells to escape surgical resection and form recurrent tumors.

**Figure 5:**
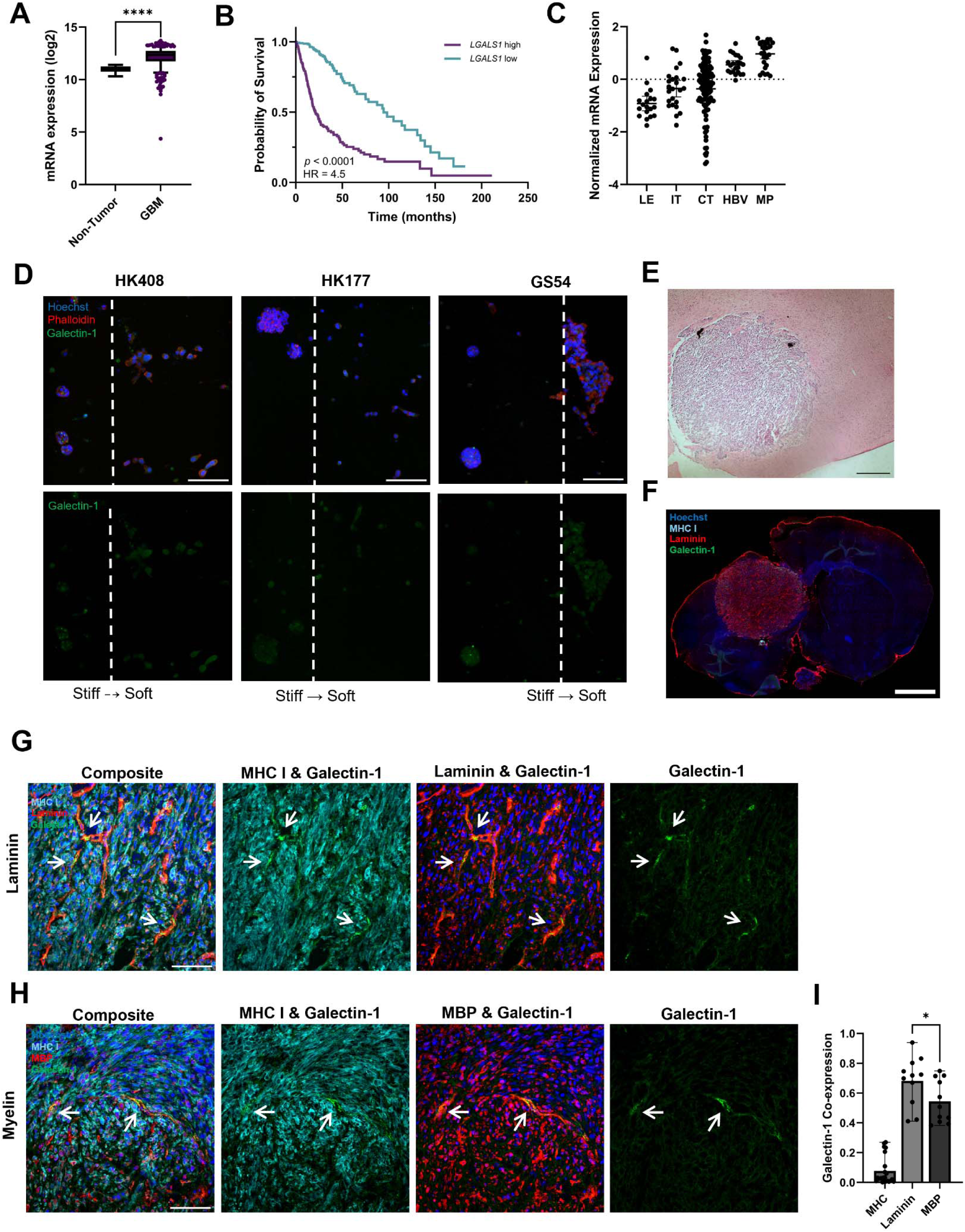
Galectin-1 is overexpressed *in vivo* GBM tissues. **A.** Data from TCGA dataset shows higher LGALS1 expression in GBM tissue compared to non-tumor tissue. Unpaired two-sided T-test, n=10 for non-tumor tissue, n = 538 for GBM tissue, \*\*\*\**p* = 3.2*10^−7^. Error bars represent 95% confidence interval. **B.** Data from TCGA dataset correlates high LGALS1 expression to a lower survival time. Log-rank, n = 333, Hazard-ratio (HR) = 4.5, *p* < 0.0001. **C.** Data from IVY GAP dataset demonstrates that galectin-1 has the highest expression along microvascular. (LE = leading edge, IT = infiltrating tumor, CT = cellular tumor, HBV = hyperplastic blood vessel, MP = microvascular proliferation). n= 19-111. **D.** ICC staining of interface hydrogels (stiff → soft). The dashed line indicates the interface. Scale bar = 100 μm. **E.**H&E staining of mouse with orthotropic xenografts of the HK408 line. Scale bar = 200 μm. **F.** Stitched full-face coronal section of the HK408 mouse xenograft brain. Section co-stained with galectin-1 (green), laminin (red) and MHC class I (HLA-ABC) to identify human tumor cells (cyan). Scale bar = 1 mm. **G.** HK408 mouse xenograft brain sections co-stained with galectin-1 (green), laminin (red) and MHC class I (HLA-ABC) to identify human tumor cells (cyan). Scale bar = 100 μm. **H.** HK408 mouse xenograft brain sections co-stained with galectin-1 (green), MBP (red) and MHC class I (HLA-ABC) to identify human tumor cells (cyan). Scale bar = 100 μm. **I**. Quantification of area of MHC+ tumor regions that are galectin-1+ (n=22) and area of laminin and myelin within MHC+ tumor regions in which galectin-1 co-stains. Unpaired two-sided T-test between laminin and myelin co-staining, n=11 images from 1 mouse, * *p* = 0.025. Error bars represent mean with range.

Immunocytochemistry (ICC) were conducted to confirm that the overexpression of *LGALS1* observed from scRNAseq resulted in expression of the galectin-1 protein. In addition to the two patient lines used in sequencing, an additional primary patient line HK408 was evaluated.^28^ Cells were cultured in the stiffness interface model (stiff → soft) for 14 days then fixed and sectioned. Sections were stained with Hoescht for cell nuclei identification, phalloidin to stain the cell cytoskeleton, and anti-galectin-1. The migrating cells express galectin-1 and have distinct morphological differences (**Figure 5D**). In the HK177 and HK408 cell lines, the non-migrating cells formed spheroids while migrating cells tended to be single cells with a more elongated morphology. In the GS54 line, migrating cells remained adjacent to the interface. However, similar morphological trends were observed for the non-migrating cells, which formed spheroids, while the migrating cells formed larger and looser clusters of cells. Cell protein lysate was harvested from the migrating and non-migrating cell populations using the Transwell interface model (stiff → soft) and 10 μg of protein lysate was loaded into each well. Protein concentration of galectin-1 was significantly higher in the migrating cells compared to the non-migrating cells in the HK177 and HK408 lines (**Figure S3**). Western blotting also confirmed that galectin-1 protein is expressed by cells in suspension culture (day 0 parent population), demonstrating that hydrogel culture is not a requirement for galectin-1 expression. The GS54 line had much lower cell migration and not enough protein could be collected from the migrating cells’ lysate to conduct a Western blot.

Next, expression of the galectin-1 protein was evaluated in mice with orthotropic xenografts of the HK408 line.^21^ GBM has been primarily found to migrate in perivascular spaces and along white matter tracts.^37^ Galectin-1 is known to interact with glycosylated structures on the neuronal cell surface, such as myelin, modulating cell adhesion to these surfaces.^38^ In sections from xenografts of the HK408 line, human tumor cells were identified with staining of MHC class I molecule HLA-ABC (MHC1). Sections were co-stained for galectin-1, MHC1, and either laminin, and a marker of vasculature basement membrane, or myelin basic-protein (MBP), to explore galectin-1 interaction with white matter tracts. Tumor was first confirmed using H&E staining in adjacent sections (**Figure 5E**). 68% (±16%) of the area in which laminin occupied tumor regions (area of overlap of laminin+ and MHC1+) co-stained with galectin-1 (area of overlap of galectin-1+, laminin+, MHC+) (**Figure 5F&G**) while 58% (±14%) of MBP+ tumor regions co-stained with galectin-1 (area of overlap of galectin-1+, MBP+, MHC+) (**Figure 5H**). Roughly 8% of tumor regions (MCH1+) had overlapping expression with galectin-1. Galectin-1 co-staining with laminin was higher than MBP (*p* = 0.025) (**Figure 5I**). Negative control side is available in **Figure S4C**.

### Inhibition of Galectin-1 reduces cell migration across the interface

To evaluate the role of galectin-1 in GBM cell migration at the mechanical interface, cells were treated with a known galectin-1 inhibitor, OTX008, which binds specifically to the amphipathic β-sheet confirmation of galectin-1, causing a conformational change that inhibits galectin-1 binding to target carbohydrates.^39, 40^ To evaluate the role of galectin-1 specifically in cell migration, the maximum concentration of OTX008 that did not affect cell viability was used. To determine the cell line-specific doses, viability was assessed 2 days in suspension culture under a range of OXT008 doses, using an ApoLive-Glo assay (**Figure S4A**). Long-term viability was confirmed through cleaved/PARP immunostaining (**Figure S4B**). The GBM lines displayed varying sensitivity to OXT008 and therefore the highest dose which did not significantly affect cell viability was chosen for each cell line which was 20 μM for HK408, 10 μM for HK177 and 1.25 μM for GS54.

GBM patient lines were encapsulated into interface hydrogels (stiff → soft) and treated every 2 days with OTX008. Cell migration was evaluated over 12 days using live cell imaging (Sartorius Incucyte S3) with images acquired every 4 hours. **Figure 6A** displays representative images for HK177 and HK408 cell migration at day 12. Cell confluency (**Figure 6B**) in the soft gel was fit to an exponential growth ODE model considering both the cell proliferation and migration rate. There was a significant reduction in cell migration in patient lines HK408 and HK177 (*p* < 0.001 and *p* < 0.01, respectively) (**Figure 6C**). The non-significant reductions in cell line GS54 may be because the viability of these cells were more sensitive to the inhibitor than the other cell lines evaluated (HK408 and HK177), which necessitated that GS54 cells were treated with a much lower concentration of OTX008. In addition, GS54 cells overall showed much less migratory activity, making it more challenging to observe changes experimentally.

**Figure 6:**
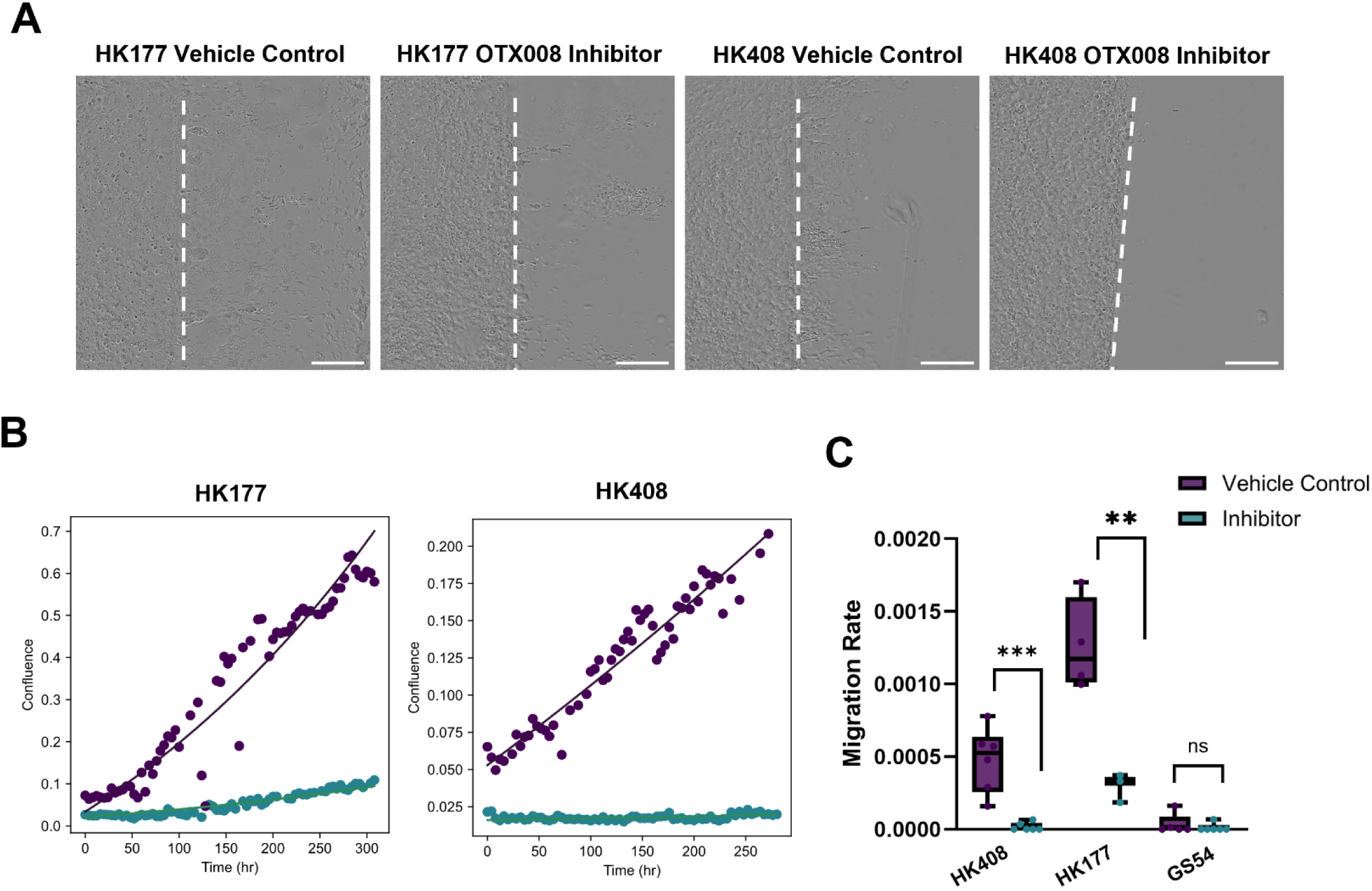
Inhibition of galectin-1 significantly reduces migration in stiff → soft interface model. **A.** Representative images of cell migration for HK177 and HK408 at the interface at d12. Scale bar = 500 μm. **B.** Cell confluence (confluence area/total area in blank gel) over course of migration experiment for representative images. Individual data points presented along with calculated growth curve. **C.** Migration rate of HK177 and HK408 is significantly lower with OTX008 inhibitor. Unpaired two-sided T-test, n = 4-6 technical replicates, ** *p* = 0.002, \*\*\**p =* 0.0003. Error bars display min and max values.

### Expression of galectin-1 is positively correlated with invasion rate in interface model

To validate that galectin-1 expression marks a pre-invasive phenotype in the HK177 line, HK177 cells were sorted based on galectin-1 expression from suspension culture. Due to the nature of galectin-1 as a glycan-binding protein, there were challenges with live-cell sorting of galectin-1, as after washing, many of the surface glycans and glycan-binding proteins were not intact, including galectin-1. To mitigate these issues, *TM4SF1*, another gene which was differentially upregulated by the invasive subpopulation in HK177, was chosen to sort invasive and non-invasive subpopulations. Expression of TM4SF1 was not present in the GS54 cell population, so this analysis was only conducted on HK177. TM4SF1+ and TM4SF1-cells were sorted using FACS (**Figure S5A**). Galectin-1 differential expression was retained over two passages, as confirmed through fixed cell flow cytometry, with 32% of cells from the TM4SF1+ group expressing galectin-1 and 9% of cells from the TM4SF1-group with measurable surface galectin-1 expression (**Figure 7A**). Pseudocolor flow plots and negative control sample available in **Figure S5B**. TM4SF1 and galectin-1 co-expression was confirmed in the xenografted mouse. In tumor regions (MHC1+), 20% of galectin-1 expression overlapped with TM4SF1 (**Figure S5C**).

**Figure 7:**
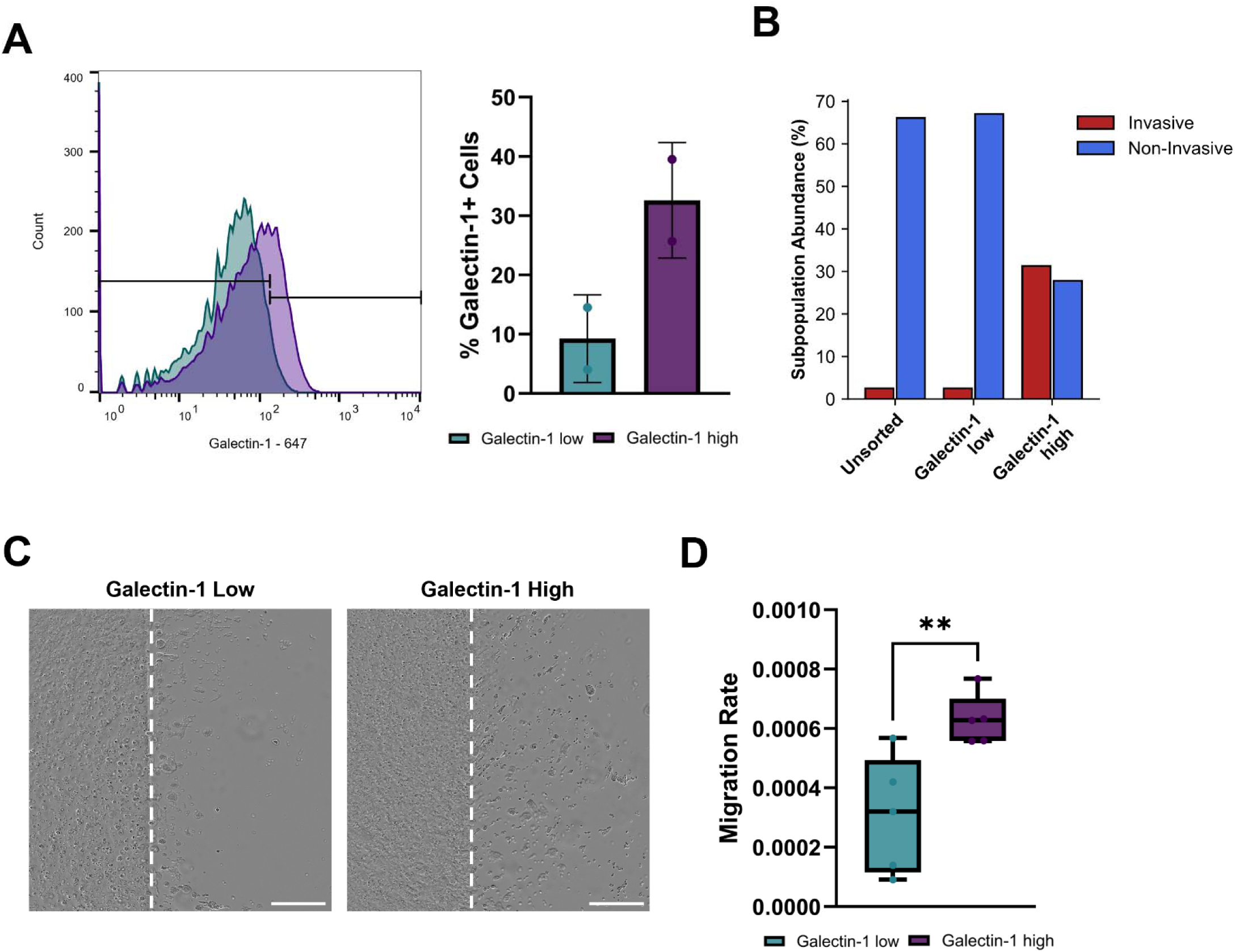
Enhanced endogenous expression of galectin-1 contributes to enhanced migration in interface hydrogel. **A.** Cells sorted based on TM4SF1 expression (high expression in purple and low expression in teal) are correlated with high and low galectin-1 expression confirmed over 2 passages (38% + vs 9%+ cells). Error bars display mean with standard deviation, n > 20,000 cells **B.** Clonal subpopulation abundance in g lectin-1 high and galectin-1 sorted populations. **C.** Representative images of cell migration for HK177 galectin-1 low and galectin-1 high groups at d10. Scale bar = 500 μm. **D.** Migration rate quantification for galectin-1 low and galectin-1 high cells. Unpaired two-sided T-test, n=5 technical replicates, ** *p* = 0.0051. Error bars dis lay min and max values.

Following the confirmation that galectin-1 was differentially expressed in the two sorted subpopulations, the subpopulation identity (invasive or non-invasive barcode sequences) was assessed in the galectin-1 high and galectin-1 low groups by targeted barcode sequencing. The unsorted parent population and the galectin-1 low group were similarly comprised and enriched for non-invasive barcode sequences (66% non-invasive, 2.5% invasive), while the galectin-1 high expressing group was enriched for the invasive barcode sequences (27% non-invasive, 31% invasive) (**Figure 7B**). The clonal identity of the 3 groups was assed 8 passages after the initial sequencing. The overall clonal diversity was reduced following extended culture where 3 invasive barcode sequences and 7 non-invasive barcode sequences were identified in the parent population and galectin-1 low group while 2 invasive and 2 non-invasive sequences were identified in flow sorted galectin-1 high group.

To confirm that the galectin-1 high and low sorted populations display different migration tendencies when separated, the migration rates of the galectin-1 high and galectin-1 low cells were evaluated once more in the interface hydrogels (stiff → soft). Evaluated at 10 days, galectin-1 high cells had a significantly higher cell migration rate (*p < 0.01*) compared to the galectin-1 low cells (2-fold change difference) (**Figure 7C/D**). These results validate that elevated galectin-1 expression in GBM cells promotes a more invasive phenotype at mechanical interfaces. Although the invasive subpopulation was a much smaller fraction of the total unsorted parent population, the invasive subpopulation could be labeled and identified by galectin-1 expression. These results further indicate that heterogeneity contributes to GBM progression through the presence of innately invasive subpopulations. Here we show that invasion is driven by a set of differentially expression genes, including galectin-1 (*LGALS1*). In this way, galectin-1 could serve as a marker of cells predisposed to migration at the tumor interface in heterogenous GBM tumors.

## DISCUSSION

Mechanical properties in the tumor microenvironment have significant effects on the invasive behavior and transcriptomic profile of GBM cells. However, due to the heterogenous nature of GBM tumors, the effect of mechanics varies across patient tumors and between subpopulations within tumors. In this study, the invasive signature of 2 patient-derived GBM lines was evaluated in a 3D culture platform mimicking the mechanical transition at the tumor-brain interface. Using this mechanically defined interface model combined with scRNAseq and lineage tracing galectin-1 was identified as a marker of invasive cells in the 2 patient-derived GBM models. While the invasion of GBM cells into the surrounding brain microenvironment is driven by many factors, here we highlight that galectin-1 mediates mechanically driven migration in GBM subpopulations. Prior studies have highlighted that mechanically sensitive cells sustain enhanced invasion and are more aggressive, leading to worse patient outcomes.^41^ Identifying the mechanically driven invasive subpopulations through high galectin-1 expression provides an opportunity to target invasive cells and potentially prolong patient survival.

The phenotypes of the two cell lines were distinct. GS54 cells were highly proliferative, less motile, and primarily expressed genes associated with the AC-like and NPC-like subtypes. In contrast, HK177 cells were more invasive and had enriched expression of genes associated with the MES-like subtype.^8^ Although the migrating cells had an increase in the number of cells with the MES-like phenotype, both the migrating and non-migrating samples from both cell lines were comprised of all cell states, thus GBM cell subtype alone is not enough to predict which cells will migrate.^9, 42^ While we were able to capture all of the malignant cell states, the cell type compositions we observed may be biased towards proliferative cells as our experimental setup required spheroid culture prior to encapsulation into the brain-mimetic hydrogel, potentially leading to loss of overall sample diversity. Nevertheless, our use of DNA barcodes allowed for the identification of distinct clonal subpopulations within the HK177 cell line, highlighting our ability to capture heterogeneity. By analyzing individual clonal subpopulations, we were able to identify transcriptomic features associated with innately invasive cells in the HK177 cell model. It should be noted that because we have defined the invasive and non-invasive subpopulations from their barcode sequence, rather than their functional migratory activity in the mechanical interface cultures, it is possible for a cell to be classified as non-migrating (from the interface culture) and also part of the invasive subpopulation (from lineage tracing). Thus, the genes identified as overexpressed in the invasive subpopulation represent genes that can be used to identify invasive subpopulations, even when not actively migrating.

By combining the interface hydrogel model and ClonMapper cell barcoding technology,^12^ distinct invasive subpopulations of HK177 which predominately migrated at the interface were identified. Among other genes, overexpression of *LGALS1* distinguished the invasive subpopulation from the non-invasive subpopulation of HK177 and *LGALS1* overexpression was a characteristic of cells which invaded across the interface in both HK177 and GS54. Galectin-1 has been reported to have multiple roles on cell behavior based on the cellular location.^43–45^ The stability of extracellular galectin-1 is dependent on its binding to glycan ligands near the cell membrane or to glycan ligands in the ECM.^43, 44, 46^ Galectin-1 preferentially binds to terminal N-linked N-Acetylglucosamine (GlcNAc) units in a β-galactoside-dependent manner with the highest binding affinity to laminin, followed by fibronectin, vitronectin, and osteopontin.^45, 47, 48^ Recently, through co-precipitation assays, galectin-1 has been identified as a binding partner for CD44, a cell surface receptor of HA, that is also upregulated in GBM.^49, 50^ The level of N-glycosylation of CD44 is correlated to cells’ ability to bind to HA, where heavily glycosylated CD44 molecules were found to inhibit cell-HA binding.^51, 52^ Previously studies have demonstrated that migration through the HA-CD44 axis is biphasic, where too little or too much HA-CD44 binding can reduce cell migration.^53, 54^ Taken together, this may suggest that galectin-1 binding to CD44 may inhibit some HA-CD44 interactions, but may enhance cell migration. However, the inhibition of HA-CD44 interactions has only been reported for galectin-9 binding to CD44.^55^

From pathway analysis, “*Asparagine N-linked glycosylation*” was upregulated in the non-migrating cells derived from the stiff matrix along with gene expression of vimentin and osteonectin. Additionally, expression of biglycan (gene *BGN*), a small proteoglycan involved in ECM assembly and an upstream signaling molecule for EGFR and IGFR signaling cascades,^56, 57^ was upregulated in the non-invasive subpopulation. Interestingly, migration of cells was not observed until roughly 5 days post-encapsulation. This delay suggests that a transcriptomic switch in response to mechanics or alterations of the ECM is required for cells to migrate. It is likely that stiffness induces the highly non-migrating cells remodel the tumor microenvironment through the deposition of glycosylated ECM which promotes the migration of the invasive subpopulation which inherently overexpress galectin-1. Remodeling of the ECM has been seen as a characteristic of GBM progression and invasion.^20, 21, 58^. Matrix mechanosensing and cell-ECM adhesion drive reorganization of the cytoskeleton through actin signaling.^59^ Previous studies have shown a link between matrix stiffness, actin signaling, and TGFβ pathway activation in serval cancers.^60–62^ Similar mechanisms may also be present in GBM tumors as we see similar pathways arise in our tumor interface model.

Galectin-1 is associated with known migrating avenues for GBM; along vasculature and white matter tracts by modulating cell adhesion to highly glycosylated structures.^37, 38, 63^ Confirmed here in an *in vivo* mouse xenograft model, galectin-1 co-stained with laminin and MPB suggesting galectin-1 aids invasive cells to migrate along vasculature and potentially along myelinated neuronal tracks in the brain.^63^ *In vivo* patient expression data of *LGALS1* in distinct spatial locations demonstrates that expression is highest along microvasculature.^64^ Microvasculature provides a physical structure and a path for GBM cells to invade from the primary tumor into surrounding tissue, with previous studies reporting that invasive GBM cells are closely associated with blood vessels.^65^ Expression of *LGALS1* by GBM cells in microvasculature niches underscores the relevance of *LGALS1* as an identifier for invasive cells across patients. A previous study suggested that the route that individual invasive GBM cells take is related to the transcriptomic state of the cells, where OPC-like and MES-like cells primarily invade along perivascular spaces while NPC-like and AC-like cells diffusely invade through the brain.^10^ Galectin-1 overexpression is associated with the MES-like phenotype.^8^ Here we present that galectin-1 identifies a unique and mechanically sensitive invasive subpopulation, identified with lineage tracing, in one patient model and marks invading cells in 3 GBM patient-models. Moreover, these invasive cells are mechanically sensitive and require galectin-1 to migrate. Linking the data presented in this study with previously reported observations, we posit MES-like GBM cells which overexpress galectin-1 have enhanced invasion along vasculature because of their ability to interact with glycosylated structures.

While galectin-1 was identified to facilitate invasion in 3 GBM patient-models, we are limited by only having the clonal subpopulation information from 1 GBM model. Additionally, the sequencing was limited to only 2 GBM patient-models, and there was a slight discrepancy in the number of detected genes and number of cells that remained following sequencing QC across the two models. These discrepancies may influence the number of identified shared differentially expressed genes between the two cell models. Although galectin-1 was not the most differentially expressed gene between the invasive subpopulations, we chose to focus on the role of galectin-1 in mediating the migration of mechanically sensitive cells as galectin-1 overexpression was also a feature of migrating cells in the HK 177 and GS54 GBM models. Other genes implicated in the migration of the invasive subpopulation of HK177, such as TM4SF1 and the metallotheians, may also be very important for the migration of mechanically sensitive invasive GBM subpopulations. Future projects will include expanding the number of barcoded patient models, sequencing additional GBM patient models, investigating additional genes implicated in GBM cell invasion, and studying the invasion of the clonal subpopulations *in vivo*. Additionally, investigating if the galectin-1 high and galectin-1 low expression subpopulations confer differential survival i*n vivo* through invasion along axons and vessels.

Using a mechanical interface hydrogel model and cellular barcodes, galectin-1 was identified as a marker of a pre-invasive phenotype and a potential target for identifying invasive GBM cells, prior to their invasion. The invasive subpopulation of HK177 was enriched by isolating galectin-1 high cells through FACS, which corresponded to higher cell invasion rates in the stiff → soft interface model. With this study, we propose that expression of galectin-1 can be used to evaluate GBM tumors for invasive capabilities and identify pre-invasive subpopulations of cells.

## METHODS

### Hydrogel Formation

Thiolated HA was prepared in-house and hydrogels as described previously.^21^ To form hydrogels, functionalized HA-SH was incorporated at a final concentration of 0.5 w/v% combined with 20kDa 4-arm PEG-SH (JenKem) and crosslinked with 20 kDa 8-arm norbornene modified PEG (JenKem) incorporated at a 1:1.2 thiol:norbornene ratio upon exposure to lithium phenyl-2,4,6-trimethylbenzoylphosphinate (LAP) (Sigma) photoinitiation (0.025 w/v%, 15 seconds of 365 nm UV exposure at 4.15 mW/cm2). RGD peptides were added to serve as cell adhesion sites, via peptide sequence GCGYGRGDSPG (GenScript) at a concentration of 0.25mM. Soft hydrogels contained a final thiol concentration of 1mM, while stiff hydrogels had a final thiol concentration of 3mM.

### Mechanical Testing

All hydrogels were swollen overnight in 1X PBS before mechanical testing. The Discovery HR20 Rheometer from TA Instruments was used for all rheological measurements. To find the shear storage modulus a frequency sweep test was performed (0.1 – 1 Hz) with 1% strain. All values are reported at 1 Hz.

### Interface Model

For migration analysis experiments, semi-circle silicon molds (5 mm x 2.5 mm) were placed into the bottom of a 48-well plate. 70 μL of non-cell containing prepolymer solution was added to the well and UV polymerized for 15s. Molds were removed and 70 μL of cell-containing prepolymer solution, at a concentration of 2000 cells/μL gel solution, were added adjacent to the cell-free hydrogel and UV polymerized for 15s. 750 μl of media was added to each well and changed every 2 days.

For sequencing experiments, 30 μL of soft hydrogel pre-cursor solution (1mM SH) was UV polymerized on the bottom of a 96-well Corning® HTS Transwell with a pore size of 8μm (Corning #3384). Cells were mixed with gel precursor solution (either 1 or 3 mM) and 30uL of cell containing solution was added to the inside of the transwell at a concentration of roughly 3000 cells/μL gel solution. Interface gels were exposed once more to UV polymerization. 300ul of media was added and changed daily.

### Cell Culture

The HK177 and HK408 patient derived lines were kindly provided by Dr. Harley Kornblum at the University of California Los Angeles. The GS54 patient derived line was kindly provided by Dr. David A. Nathanson at the University of California Los Angeles. GBM cells were cultured in DMEM/F12 media (StemCell Technologies #36254) containing 20 ng/mL human FGF (PeproTech #100-18B), 50 ng/mL human EGF (PreproTech #AF-100-15), 5 ng/mL heparin (Sigma #H3149-100KU), 2% Gem21 NeuroPlex supplement (GeminiBio #400-160) and 2 mmoL of L-Glutamine (StemCell Technologies #07100), and 100 μg/mL Normocin (Invitrogen #ANTNR1) at 37C 5% O2.

### Flow Cytometry

For barcoding cell identification cells were collected and passaged with 1X TrypLE (Thermo Fisher #12604021) for 5 minutes. Cells were pelleted and washed 2X in cold PBS (Fisher Scientific # BP2944100). Cell pellets were resuspended in cold PBS stained with 7-AAD (7-Aminoactinomycin D) (Invitrogen A1310) according to the manufacturer’s recommendations for dead cell identification. Cells were kept on ice until sorting. For sorting of TM4SF1/Galectin-1 high expressing cells, cells were passaged with 0.5mM EDTA-PBS (EDTA Sigma #E9884) with gentle agitation for 5 minures to preserve surface proteins. Cells were pelleted and washed 2X in cold 1%BSA-PBS (BSA #9048-46-8) Cells were sorted on the Sony SH800S Cell sorter using a 100 μm microfluidic sorting chip. Following cell sorting, cells were expanded until cell numbers reached a minimum of 3*10^6^ requiring 5 passages for the barcoded cells and 3 passages for the galectin-1 high and low expressing population. Following expansion cells were cryopreserved until use in experiments.

For galectin-1 and TM4SF1 surface staining fixed cell studies, cells were fixed with 4% PFA for 15 minutes and then washed with 1%BSA-PBS. All samples were incubated on ice for 30 minutes with primary antibody; rabbit anti-TM4SF1 (1:50, Invitrogen PA5-21119) and mouse anti-galectin-1 (1:25, Santa Cruz Biotechnology sc - 166618). Following incubation cells were pelleted and washed with 1%BSA-PBS. Cells were incubated for 30 minutes at 4C with secondary antibody at a 1:100 dilution (in their respectively secondary antibodies at a 1:500 dilution (anti-rabbit Alexa Fluor 488 and anti-mouse Alexa Fluor 647). Cells were washed 2X with 1%BSA-PBS. Cells were resuspended in cold PBS and kept on ice until analysis. A negative control sample (only secondary antibody) was used to determine positively stained cells. Cells were run on a Attune™ NxT Flow Cytometer.

### Cell Barcoding

Cells were barcoded according to the ClonMapper protocol as described previously.^13^ Briefly, cells were transduced with barcode-containing lentivirus at a MOI of 0.1. 72 hours post transduction, BFP+ barcoded cells were sorted using FACS. (**Figure S1A**).

Assessment of barcode diversity in samples was conducted according to the ClonMapper protocol as described previously.^13^ Briefly, DNA was extracted using DNeasy Blood & Tissue Kit (Qiagen #69504) per manufacturer’s recommendations. Genomic DNA was processed for next-generation sequencing (NGS) in a two-stage PCR reaction. In stage 1, barcode sequences were amplified, and in stage 2, Illumina adapters and indices were ligated to barcode sequences for their identification following NGS. Barcode libraries were sequenced at UT Austin’s Genomic Sequencing and Analysis Facility (GSAF). Cells were sequenced on an Illumina NovaSeq S1 with 100 cycles with a minimum of 5 million reads per sample. To quantify barcode abundance from targeted barcode sequencing, Pychasier was used with default parameters.^66^

### ScRNAseq Cell Preparation

One scRNAseq run was conducted for the barcoded HK 177 cells and a separate run for the GS54 line. For each cell line, 30 hydrogels of each from each condition (soft → soft migrating, soft → soft non-migrating, stiff → soft migrating, stiff → soft non-migrating) were pooled together prior to cell extraction. Once pooled, hydrogels were dissociated and cells were extracted using the human tumor kit and the GentleMACs human tumor dissociator from Miltenyi Biotec (130-095-929). Cells were washed 3X with 1X PBS and stained with 7AAD according to the manufacturer’s recommendations for dead cell identification. Cells were sorted using FACS to remove any gel particles and to ensure that only live cells were submitted for sequencing. Prepared cells were submitted to UT GSAF for sequencing preparation. Libraries were prepared using 10X Genomics’ Chromium Single Cell 3 v3 kit. 1000 cells per condition of the HK177 line and 1500 cells per condition of the GS54 line were sequenced on an Illumina NovaSeq S1 with 100 cycles at a depth of 50,000 reads/cell.

### ScRNAseq Data Analysis

Annotated count matrices were generated from the raw fastq files using Cell Ranger version 3.0.2. Genes were aligned to the reference genome NIH GRCh38. Quality control was conducted on the number of counts per cell, number of genes per cell, and the fraction of counts from mitochondrial genes as described by Heumos, et al.^67^ Low quality data was removed from the dataset using a median absolute deviation (MAD) threshold of 3 and a mitochondrial gene fraction > 0.2. Doublets were detected and removed using Scrublet, using an expected doublet rate of 2.3%. Genes that were not identified in at least 10% of cells were removed from the dataset. In the GS54 line, 4943 cells and 9619 met the QC cutoffs. 1519 cells in the soft → soft migrating, 1331 cells in the soft → soft non-migrating, 1157 cells in the stiff → soft migrating, and 936 cells in the stiff → soft non-migrating. In the HK177 line, 3850 cells and 8873 genes met the QC cutoffs. 786 cells in soft → soft migrating, 1143 cells in soft → soft non-migrating, 998 cells in the stiff → soft migrating, and 923 cells in the stiff → soft non-migrating. For visualization, cells were assigned a cell cycle score based on expression of cell cycle canonical markers and regression was performed to remove individual variances due to cell cycle.^68^ For differential expression and expression quantification, data was normalized as log(CPM+1), where CPM is counts per million.

Following QC, NGS reads from Cell Ranger were processed with Pycashier to extract the UMI, cell barcode and ClonMapper DNA barcode.^13^ Duplicate reads (same cell barcode and UMI) were removed. Due to amplification and sequencing errors, some ClonMapper barcodes were read with aberrant bases or truncated sequences. To correct for this, short lineage reads were aligned to longer reads to assign them to a full 20-nt barcode. To correct misreads during sequencing, lineages with less than 3 base pair differences across the 20-nt barcode were merged and assigned to the same barcode. Cells which had more than one barcode sequence were assigned to the barcode with higher UMI counts. Finally, barcodes which had a UMI count of 5 or less were removed. Cells that we were unable to confidently assign a lineage to were assigned ‘none’ and not used in lineage analysis but were used in differential gene analysis.

Subsets of the data aggregated by clonal subpopulation or by sample identity (substrate stiffness and migration status) underwent differential gene analysis was conducted using MAST as described previously.^69^ To control for covariance in experimental conditions, when not being actively tested, barcode, substrate stiffness, and migrating status were included as covariates in MAST. For the analysis of differential gene analysis across the two cell lines, the cell line was used as an additional covariate. False discovery rate control was performed using Benjamini-Hochberg-adjusted p-values. Downstream analysis was conducted on genes with an adjusted p-value of *p* < 5*10^−5^. Pathway analysis was conducted using Ingenuity pathway analysis (IPA, Qiagen).

For cell subtyping, the cells were assigned a cell state based on expression of marker genes as identified and described by Neftel, et al.^8^ MES 1 and MES 2 markers were combined into a single MES subtype and with NPC 1 and NPC 2 were combined into a single NPC subtype. Not all of the Neftel marker genes were observed in each cell line so a subset of the marker genes that were expressed by cells were used to score and assign cell types. The genes for each subtyping analysis can be found in the supplemental data. A two-sided unpaired t-test was used to calculate differences in MES, AC, OPC, and NPC score per cell between conditions. False discovery rate control was performed using Benjamini-Hochberg-adjusted p-values.

### Animal Model

Sections of mice with orthotropic xenografts of the HK408 line were taken from previous experiments.^21^ Briefly, 50,000 cells were injected into the brains of Nod-SCID-gamma (NSG^TM^) (Strain #:005557) mice. Tumor growth was monitored using VIS imaging and weighed weekly. Mice were monitored for wellbeing and were euthanized when mouse weight reached 80% of initial weight. Mice were perfused with 4% paraformaldehyde (PFA) and brains were harvested for downstream analysis.

### Mouse Brain and Hydrogel Sectioning

Mouse tissue and hydrogels were fixed with 4% PFA for 48 hours (mouse) and 1 hour (hydrogels). Following fixation samples were washed thrice with 1X PBS. For embedding, samples were incubated in 5% sucrose-PBS at room temperature for 1 hour, followed by 48-hour (mouse) or overnight (hydrogels) 4°C incubation in 30% sucrose-PBS. Mouse tissue was embedded in 30% sucrose-OCT while hydrogels were embedded in 20% sucrose-OCT for and incubated for 3 hours before being flash frozen. Embedded hydrogels and mouse tissue was stored in −80°C prior to cryosectioning. Preserved hydrogels and tissue were sectioned at a 14 μm thickness. Mouse brains were sectioned coronally at 14 μm thickness. Sections were kept at −20°C until staining.

### Immunocytochemistry

The sections were air-dried for approximately 10 minutes before fixing in 4% PFA for 12 minutes. Samples were rinsed thrice in 1X TBS for 5 minutes each. Tissue sections were blocked in 2% BSA and mouse-on-mouse blocking, according to manufacturer’s recommendations (Vector Labs, MKB-2213-1) in 1X TBS for 1 hour. Hydrogel sections were blocked in 4%BSA-TBS for 1 hour. Mouse sections were incubated overnight in blocking solution at 4C with the following antibodies: chicken anti-MBP (1:100, Invitrogen PA1-10008), rat anti-HLA-ABC (1:100, Invitrogen MA1-80014), rabbit anti-laminin (1:100, Invitrogen PA1-16730), rabbit anti-TM4SF1 (1:100, Invitrogen PA5-21119), and mouse anti-galectin-1 (1:50, Santa Cruz Biotechnology sc - 166618). Hydrogel sections were incubated in mouse anti-galectin-1 (1:50). The following day slides were rinsed thrice in 1X TBS for 5 minutes. For OTX008 inhibitor studies slides were incubated in Cleaved PARP primary antibody solution at a 1:400 dilution (Cell Signaling 5625). Slides were incubated for 1 hour at room temperature in their respectively secondary antibodies at a 1:500 dilution (Alexa Flour 488, 555 and 647, Invitrogen) along with Hoechst 33342 (ThermoFisher Scientific 62249) for nuclear labeling at a 1:500 dilution. Following incubation slides were rinsed thrice in 1X TBS for 5 minutes. Slides were mounted and stored at 4C until imaging. Fluorescence images were taken using the Leica SP8 laser confocal microscope, 10× and 20× objective lenses.

Mouse section image analysis and quantification were performed in python. Individual channels were separated and fluoresce intensity was thresholder based on negative control slides. Tumor regions were determined by summing the area of MHC1+ pixels above the threshold value. For quantification of laminin, MBP and galectin-1 stains, only areas where stained pixels were above the threshold value and overlapped with MHC1+ pixels were used. Galectin-1 co-staining area was calculated as area of overlapping pixels with laminin or MBP in tumor regions per total galectin-1 staining area.

### Western Blot

Cells were cultured in interface conditions for 2 weeks. Two biological replicates were conducted and for each cell line, and 2 separate western blots were run for each biological replicate. For each cell line in each replicate, 12 hydrogels of each from each condition (stiff → soft migrating and stiff → soft non-migrating) were pooled together prior to protein lysate extraction. Hydrogels and 2*10^6^ cells from suspension culture were collected into tissue homogenizer tubes (Qiagen, 13114-50) containing 200uL of Ripa lysis buffer (Thermo Fisher, P189901) with 2 uL of Protease inhibitor cocktail (Cell Signaling, 5872S). Samples were homogenized in a Bead Mill 24 Homogenizer (Fisher Scientific, 15-340-163). Lysate was collected and moved to 1.5 mL microcentrifuge tubes centrifuged at 21,100 xg for 30 minutes at 4C. Surprenant was collected and flited through Ultrafree-MC centrifugal units (Sigma, UFC40GV25) and centrifuged at 12,000 xg for 4 minutes at at 4C. Flow through was collected, aliquot and stored at –80C until further use. Protein concentration was determined using a microBCA assay (Thermo Fisher, 23235). 10 μg of protein lysate was loaded into each well of a 4-20% Tris-glycine protein gel (Thermo Fisher, XP04202BOX) and run according to the manufacturer’s instructions. Following transfer, blots were blocked in 4% BSA-TBS for 1 hour at room temperature followed by overnight primary antibody incubation (4C) with mouse anti-galectin-1 (1:500, Santa Cruz Biotechnology, sc - 166618) and rabbit anti-GAPDH (1:1000, Cell signaling, D16H11). The following day blots were washed 3 x 5 minutes in TBST before 1 hour of incubation at room temperature with rabbit HRP-conjugated antibody (1:2000, Cell Signaling, 7074S) and mouse HRP-conjugated (1:2000, Santa Cruz Biotechnology, sc-525416). Following incubation blots were washed 3 x 5 minutes in TBST beofre 5 minutes of incubation with developing buffer (Thermo Fisher, 34580).

### OXT008 Inhibitor Study

OTX008 (MedChemExpress, MCE HY-19756) was prepared at stock concentration of 5mM in DMSO, aliquoted and stored at −80C. To determine the cell line specific doses, doses curves from 20uM to 0.625 uM were applied to each cell line in triplicates. Viability was assessed using ApoLive-Glo™ Multiplex Assay (Promega, G6410). For migration studies, cells were encapsulated and treated with the cell-line specific doses. Media was changed with fresh OXT008 every 2 days for the duration of the experiment.

Long-term viability was confirmed through cleaved/PARP immunostaining. Following the duration of the experiment, hydrogels were fixed, sectioned, and stained as described above. Viability was determined by counting the total live nuclei (non-stained) divided by the total number of nuclei.

### Migration Imaging and Rate Calculation

For migration analysis studies, interface hydrogels were imaged using Incucyte S3 (Sartorius) with the 4X objective using phase contrast imaging. Images were taken every 4 hours for the duration of the experiments. Cell confluence masks were generated at each time point from images with Incucyte built-in default segmentation. Images that deviated from the median focus position by more than 10% were removed from the analysis. To control for variation in camera position, each image was aligned to the image from the previous timepoint. To align images, an optimal x,y position was iteratively calculated that maximized the overlap in confluence masks of the two images. The location of the gel interface was computationally determined from the confluence mask of the first (t=0h) image by maximizing the number of confluent pixels to the left of the interface position and minimizing the number of confluent pixels to the right of the interface position. New cells in the test gel were quantified as the percent confluence to the right of the interface over time. Time-series confluence data in the test gel was fit to a standard exponential growth ODE with an added migration rate m to model new confluency from migrating cells:

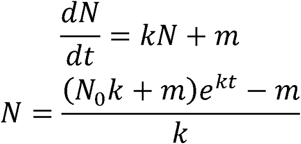

Here, *N* is the number of cells (confluence), *k* is the proliferation rate (1/hour), and *m* is the migration rate (confluence/hour). Optimal parameters for k and m were individually fit to each replicate by Levenberg-Marquardt optimization. Significance of differences in growth rate and migration rate between conditions were tested by an unpaired two-tailed t-test on fitted parameter estimates.

### Statistics and Reproducibility

Statistical analyses are described in individual sections and in the figure captions. For scRNaseq differential gene analysis, *p* < 5*10^−5^ was used to identify significantly different genes. The number of samples, n, refers to distinct samples in which measurements were taken. For samples with two conditions, data were analyzed using an unpaired two-tailed t-test and for >2 samples an ordinary one-way Analysis of Variance (ANOVA) was conducted, followed by Tukey’s post hoc analysis using GraphPad Prism 10. Statistical significance was denoted as follows: not significant (ns, p > 0.05), \**p* < 0.05, \*\**p* < 0.01, \*\*\**p* < 0.001, and \*\*\*\**p* < 0.0001. +/− in figures and results represent the standard deviation.

## Supporting information

Supplemental Data File 1 which contains the genes used to determine the GBM malignant cell type

Supplemental Data File 2: The statistical analysis for determining if the malignant cell type composition is significantly different between each cond

## Acknowledgment

This work was supported by the National Institutes of Health NCI R01CA241927-01A1 (SKS, AB) and U01CA25354001 (AB), the Cancer Prevention and Research Institute of Texas RR210042 (SKS), and the American Brain Tumor Association (DG1800018, SKS).

Next generation sequencing was performed by the Genomic Sequencing and Analysis Facility at UT Austin, Center for Biomedical Research Support (RRID#: SCR_021713).

Florescence activated cell sorting was conducted at the Microscopy and Flow Cytometry Facility at UT Austin (RRID:SCR_021756).

The computing for this project was performed at the University of Texas at Austin Biomedical Research Computing Core Facility (RRID:SCR_021979).

The HK177 and HK408 patient derived lines were kindly provided by Dr. Harley Kornblum at the University of California Los Angeles. The GS54 patient derived line was kindly provided by Dr. David A. Nathanson at the University of California Los Angeles.

We would like to thank Mollie J Harrison and Ayla Farzamnia for their assistance in acquiring additional images.

## Author contributions

Conceptualization TS, SKS; Data curation, TS; Formal analysis, TS, MC; Funding acquisition, SKS, AB; Investigation, TS, MC, NA; Methodology, TS, MC; Project administration, SKS; Resources, TS, MC, SKS, AB; Software, TS, MC; Supervision, SKS, AB; Validation, TS; Visualization, TS; Writing – original draft, TS; Writing – review & editing, TS, MC, SKS, AB.

## Data Availability

The raw fastq and processed count matrices from scRNAseq discussed in this publication have been deposited in NCBI’s Gene Expression Omnibus (Edgar *et al*., 2002) and are accessible through GEO Series accession number GSE328755 (https://www.ncbi.nlm.nih.gov/geo/query/acc.cgi?acc=GSE328755). The source data for the figures can be accessed on the Texas Data Repository at https://dataverse.tdl.org/dataverse/interface-invasion.

## Code Availability

The code used to generate data is available at https://doi.org/10.5281/zenodo.23512182.1 Cell Ranger version 3.0.2 was used to generate count matrices from scRNAseq.

## Ethics approval statement

Institutional Biosafety Committee approval IBC-2025-00057.

