## Supplemental Figures S1-S5 for "Galectin-1 identifies a unique subpopulation of highly invasive glioblastoma cells and enables their migration"

**Supplemental Figure 1: ClonMapper cell barcodes identified in HK177 sample**

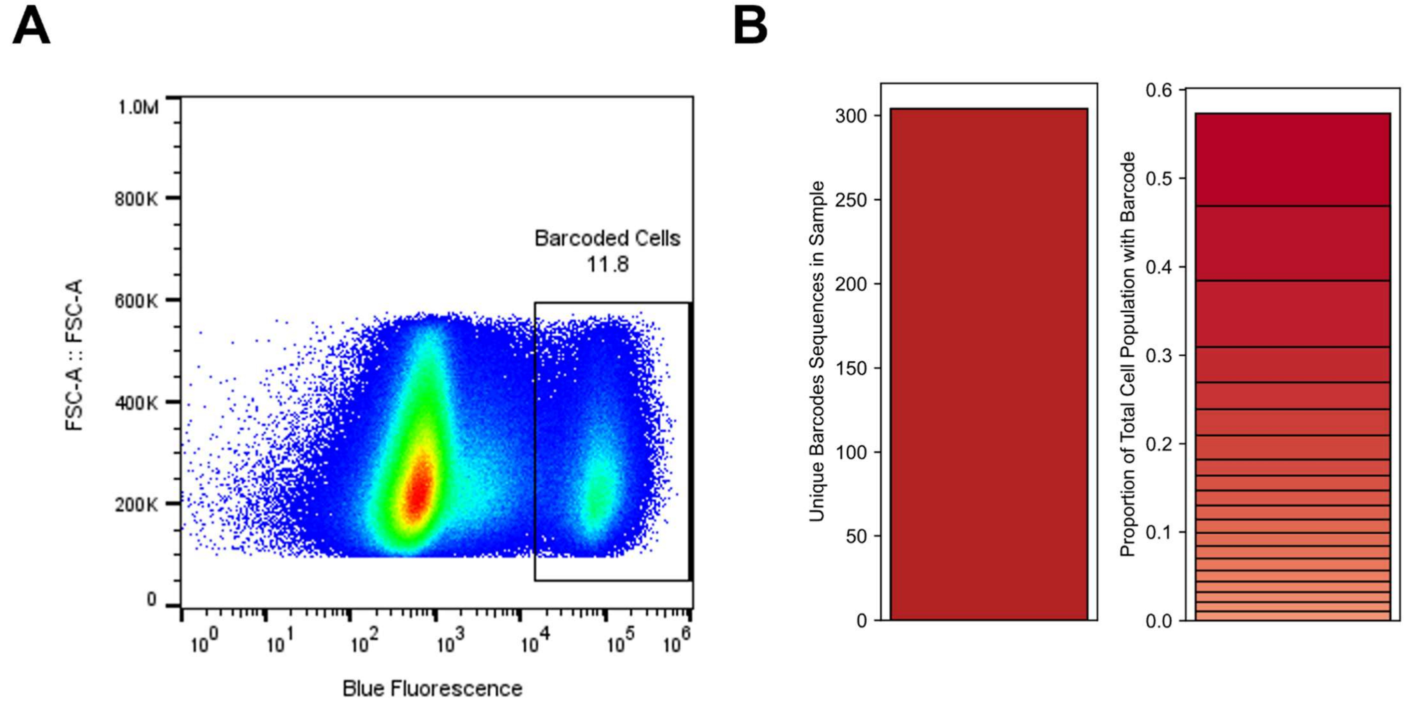


**Figure S1:** ClonMapper cell barcodes identified in HK177 sample. A. Flow Cytometry confirms barcoded cells via bfp expression. B. Number of unique barcode sequences identified in sample and and abundance of the 20 highest expressed barcode sequences.

**Figure S2: Characteristics of the invasive and non-invasive subpopulation of HK177**

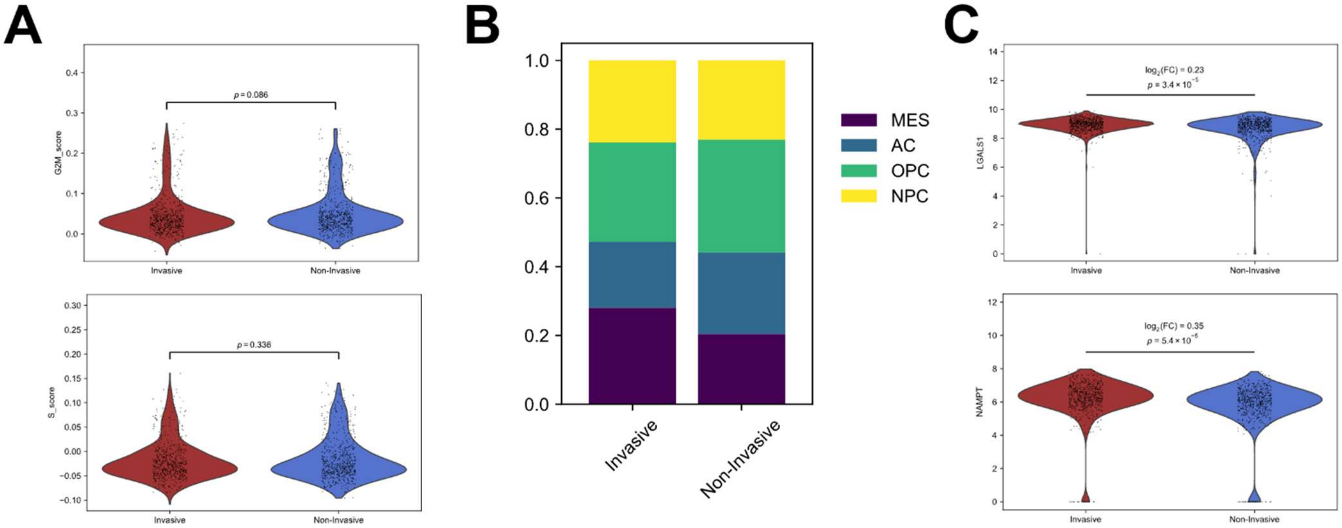


**Figure S2:** Characteristics of the invasive and non-invasive subpopulation of HK177. **A.** Cell phase score compared between the invasive and non-invasive subpopulation. Unpaired two-sided T-test between all cells with an invasive barcode sequence (n = 641) and all cells with a non-invasive barcode sequence (n = 655). No significance. **B.** Malignant cell type characterization of the invasive and non-invasive subpopulations. The full states table for all comparisons of both cell lines can be found in Table S2. Unpaired two-sided t-test. **C.** LGALS1 and NAMPT are identified from MAST as differentially expressed between subpopulations when controlling for cell cycle phase and MES cell type score.

**Figure S3: Protein expression of gelactin-1 from *in vitro* interface model**

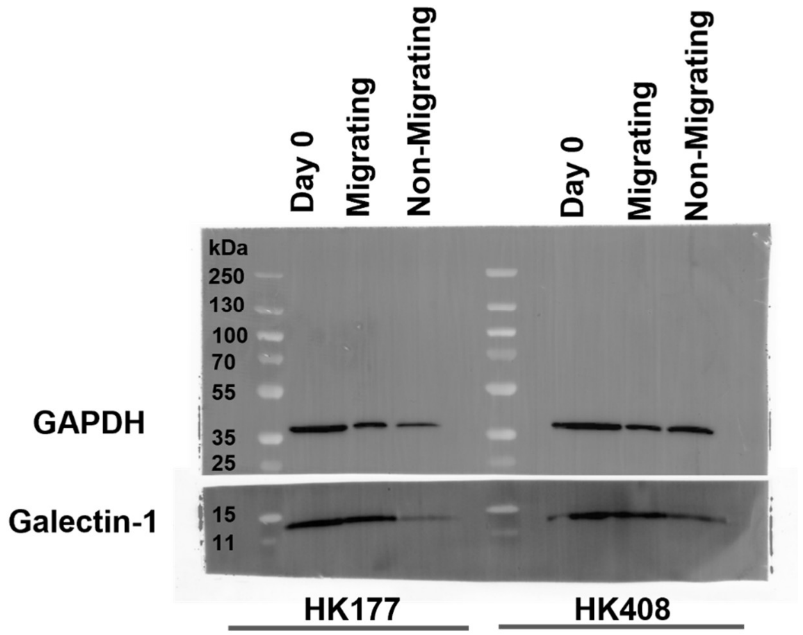


**Figure S3:** Western blot confirms that there is higher galectin-1 protein expression in migratory cells compared to the non-migratory cells in HK177 and HK408 in stiff → soft interface hydrogel. 10 µg of protein lysate was loaded into each well.

**Figure S4: Long term use of OTX008 is not cytotoxic at cell line-specific does**

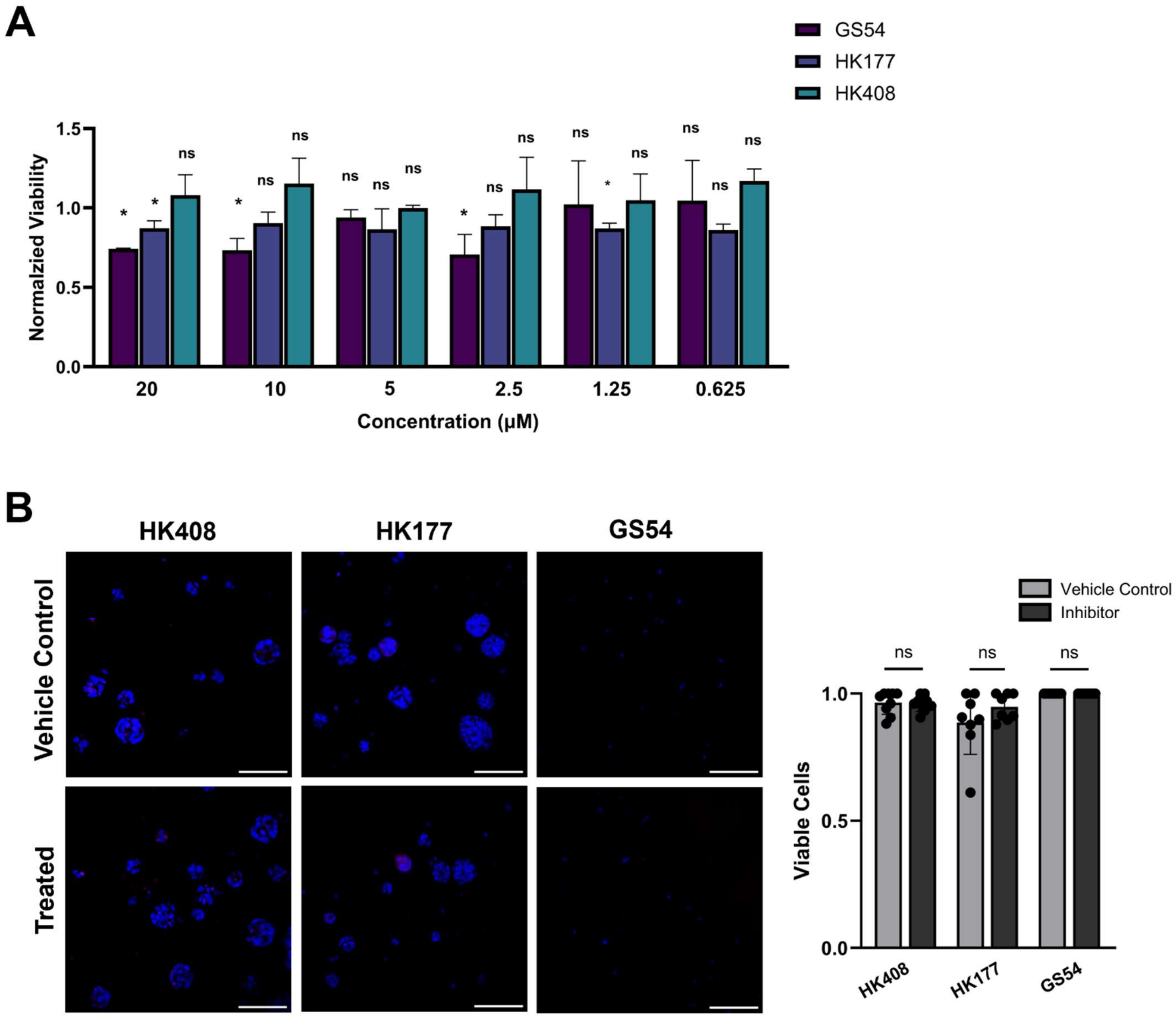


**Figure S4:** Treatment with OXT008 is not cytotoxic at cell line-specific does**. A.** ApoGlo Live viability assay to determine concentration that does not reduce cell viability in cell culture condition. One-way unpaired t-test, n=3 technical repeats * $p$ < 0.05, ns= no significance. **B**. Long term viability determined in stiff hydrogels after 12 days in culture. Cells stained with Hoescht nuclear staining (blue) and cleaved/parp to stain for dead cells (red). Scale bar = 100 µm. Unpaired two-sided T-test, n=8, ns = no significance.

**Figure S5: TM4SF1 and Galectin-1 co-expressed by HK177 cells and in *in vivo* xenograft xenografts of GBM line HK408.**

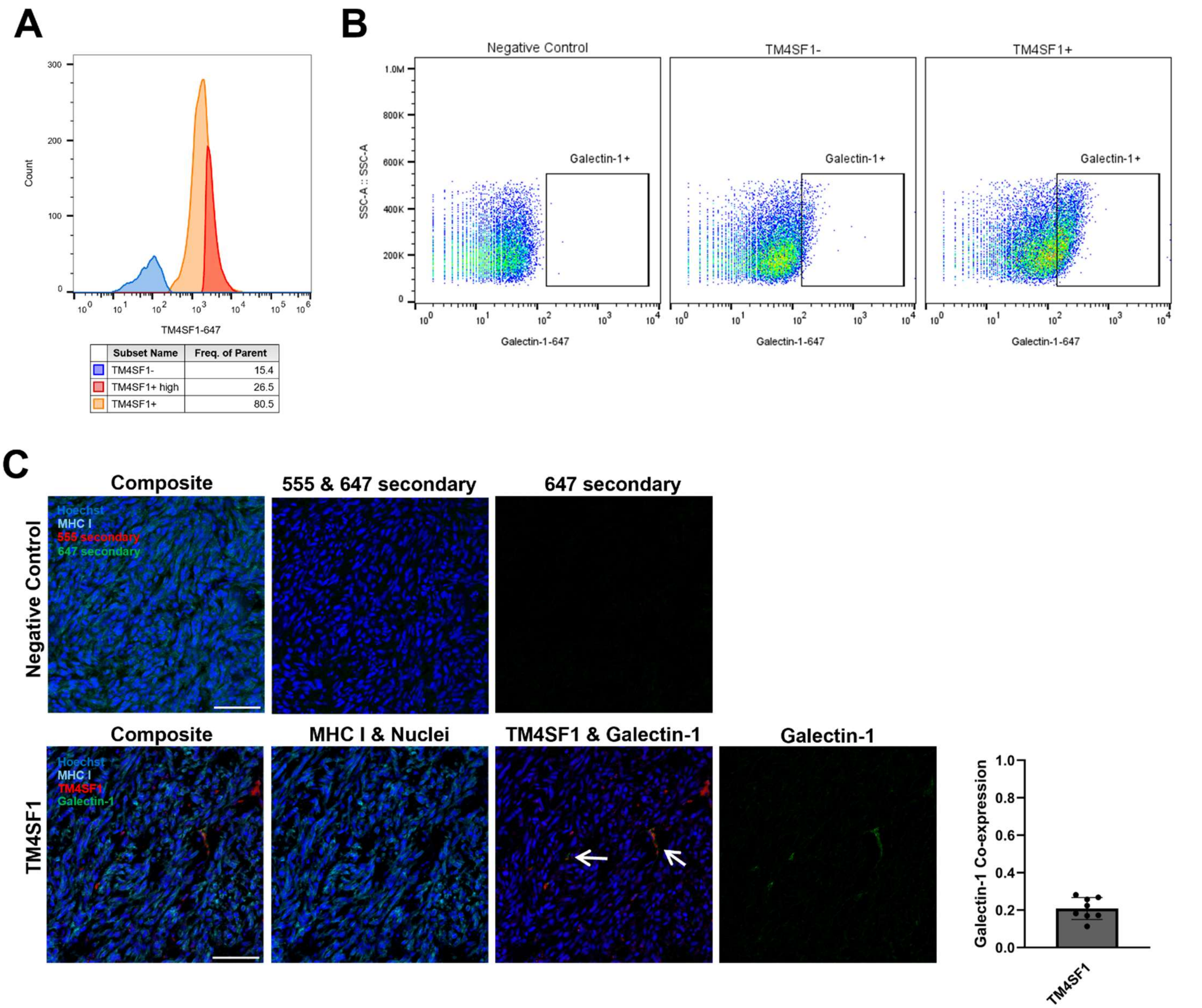


**Figure S5.** TM4SF1 and Galectin-1 co-expressed by HK177 cells and in *in vivo* xenograft xenografts of GBM line HK408. **A.** TM4SF1+ high expressing cells (red) and TM4SF1- (blue) cells were sorted using FACS. **B.** Pseudocolor plots demonstrate that TM4SF1+/- sample corresponds to high and low galectin-1 expression. **C**. Negative control for ICC staining of orthotropic xenografts. Scale bar = 100 μm. HK408 mouse xenograft brain sections co-stained with galectin-1 (green), TM4SF1 (red) and MHC class I (HLA-ABC) to identify human tumor cells (cyan). Scale bar = 100 μm. Quantification of area of co-stain on right of panel. Error bars represent mean with standard deviation.
